# Host traits impact the outcome of metagenomic library preparation from dental calculus samples across diverse mammals

**DOI:** 10.1101/2025.03.19.643754

**Authors:** Markella Moraitou, Johnny Richards, Chanah Bolyos, Konstantina Saliari, Emmanuel Gilissen, Zena Timmons, Andrew C. Kitchener, Olivier S. G. Pauwels, Richard Sabin, Phaedra Kokkini, Roberto Portela Miguez, Katerina Guschanski

## Abstract

Dental calculus metagenomics has emerged as a valuable tool for studying the oral microbiomes of humans and a few select mammals. With increasing interest in wild animal microbiomes, it is important to understand how widely this material can be used across the mammalian tree of life, refine the related protocols and understand the expected outcomes and potential challenges of dental calculus sample processing. In this study we significantly expand the breadth of studied host species, analysing laboratory and bioinformatics metadata of dental calculus samples from 32 ecologically and phylogenetically diverse mammals. While we confirm the presence of an oral microbiome signature in the metagenomes of all studied mammals, the fraction recognised as oral varies between host species, possibly due to both biological differences and methodological biases. The overall success rate of dental calculus processing, from extractions to sequencing, was ~74%. Although input sample weight was positively associated with the number of produced library molecules, we identify a negative impact of enzymatic inhibition on the library preparation protocol. The inhibition was most prevalent in herbivores and frugivores and is likely diet-derived. In contrast, hosts with an animalivore diet posed fewer challenges during laboratory processing, and yielded more DNA relative to sample weight. Our results translate into recommendations for future studies of dental calculus metagenomics from a variety of host species, identifying required sample amounts, and emphasising the utility of dental calculus in exploring the oral microbiome in relation to broader ecological and evolutionary questions.

## Introduction

Host-associated microbiomes contribute to a multitude of fundamental biological functions of their hosts. The seemingly endless list includes processes as diverse as immune response (Ippolito et al. 2018, 202), reproduction (Comizzoli et al. 2021), nutrient metabolism (Rowland et al. 2018) and thermoregulation (Chevalier et al. 2015). As most studies focus on the human gut, the microbiomes of non-human animals, especially of body sites other than the gut, remain largely understudied. However, it is unclear how far the knowledge from the human gut microbiome extends to the wider biological diversity of hosts and microbial communities. Animal microbiomes can have important implications for conservation and adaptation (Trevelline et al. 2019; Alberdi et al. 2016), act as reservoirs for zoonotic diseases (White and Razgour 2020) and respond to anthropogenic impacts, such as habitat degradation (Barelli et al. 2015; Amato et al. 2019) and contamination (Lapanje et al. 2010; Costa et al. 2016; Brealey et al. 2021).

While less well studied than the gut microbiome, the oral microbiome has systemic effects on its host: it is involved in a diversity of systemic diseases (Nakano et al. 2009; Teles and Wang 2011; Olsen and Singhrao 2015; Flynn, Baxter, and Schloss 2016; Gao et al. 2016; Warinner 2016) and plays an important role in the production of nitric oxide (Duncan et al. 1995), a molecule that participates in blood flow regulation and inflammation (Ghimire et al. 2017). At the same time the oral cavity is more exposed to the external environment than the gut microbiome and likely contains environmental microbes (Shaiber et al. 2020), which may offer additional insights into the host’s habitat. Yet, collecting samples from wild animals remains challenging due to hurdles both logistic (fieldwork in remote locations, need for research permits, etc.) and ethical (potentially affecting the health and welfare of rare or endangered species). This is particularly true for microbiomes other than the gut, as faecal samples can often be easily obtained without disturbance to the host individuals, in contrast to sampling from other body sites, which may require handling. Studying captive animals may lead to biassed inferences, as, at least for the gut, captivity was shown to drastically alter and homogenise the microbiome (McKenzie et al. 2017; Clayton et al. 2016).

Dental calculus, the mineralised form of the dental plaque microbiome, offers a way to overcome many of the challenges presented above. Dental calculus forms on the surface of teeth through periodic mineralisations (Schroeder and Shanley 1969; Jepsen et al. 2011) of the microbial biofilm, entrapping and preserving the DNA of the oral microbiome, the host, and dietary components (Mann et al. 2018; 2020; Warinner et al. 2015). It is widespread among mammals and readily available from museum specimens (Brealey et al. 2020, Richards et al. in prep) and archeological material (Warinner et al. 2014; Armitage 1975; Ciochon, Piperno, and Thompson 1990; Dobney and Brothwell 1987), which allows researchers to circumvent the logistic and ethical difficulties of handling wild animals. In addition, due to the exceptional preservation of genetic material (Velsko et al. 2019; Fellows Yates et al. 2021), this ‘microbial fossil’ can be sampled decades, centuries, or even millennia after the host’s death, and analysed using metagenomics, providing snapshots of past microbiomes (Warinner et al. 2014). This has allowed for temporal studies of the oral microbiome, helping to understand changes in microbiome structure and function in the distant past (Quagliariello et al. 2022; Velsko et al. 2024; Gancz et al. 2023) or more recently, under increasing human impact on the environment (Brealey et al. 2021). Beyond its primary use for microbiome research, dental calculus can be used for host genomics. Because the eukaryotic content is low in this material (<1%; Mann et al. 2018), shotgun metagenomics provide only a limited view of the host, but may still allow for molecular sexing, mitochondrial DNA assemblies, and low coverage host genomics (Brealey et al. 2020; Moraitou et al. 2022; Gower et al. 2019). Alternatively, researchers may opt for target enrichment techniques to increase host content (Ozga et al. 2016; Ziesemer et al. 2019).

Despite all these possible applications, dental calculus has so far been predominantly used to study the oral microbiome of humans (see Fotakis et al. 2020; Velsko et al. 2024; Gancz et al. 2023; Warinner 2016; Granehäll et al. 2021; Klapper et al. 2023, among others) and a small number of non-human species (Moraitou et al. 2022; Brealey et al. 2021; 2020; Ozga and Ottoni 2021; Ottoni et al. 2019). Given its broad presence across a diverse set of mammals and the wide scope of questions in ecology and evolution that can be addressed with this material, we can expect its increased use in the future. Therefore, we need to better understand which factors determine the success of molecular analyses based on dental calculus. While its sampling is usually less damaging to a specimen than sampling dentine or bone, it is still considered destructive, as the sampled material is used up during laboratory processing. Although dental calculus reportedly preserves DNA better than dentine (Mann et al. 2018), the sample success rate is around 70-80%, with some evidence of sample inhibition (personal observations). Therefore, to minimise unnecessary sampling of rare museum specimens, and to better manage project resources and funding, it is crucial to understand factors that influence success rate in the laboratory and hence identify best sampling strategies.

This study aims to assess the efficiency and success rate of Illumina metagenomic library preparation from dental calculus, depending on the characteristics of the collected samples and host species. In the absence of any inhibition we expect that sample weight will be positively correlated to DNA concentration in a standardised extraction and to the quantity of fragments that are successfully turned into next generation sequencing libraries. However, we hypothesise that additional factors, such as host diet and/or taxonomic order, may impact the outcome of these processes. Regarding host diet, plants can contain secondary compounds, such as polyphenols, polysaccharides, tannins, etc., that inhibit enzymatic reactions (Schrader et al. 2012, 202). These are often co-extracted during sample processing, generating brown or yellow DNA extracts (Stevenson 1994). Therefore, we expect that dental calculus from herbivorous species with high content of such compounds in their diet will exhibit enzymatic inhibition, negatively affecting laboratory protocols. Regarding host taxonomy, dental calculus from phylogenetically and ecologically diverse host species may differ in DNA yield as a result of structural differences in dental calculus, for instance related to deposit thickness or texture, which may have a phylogenetic signal. To test these hypotheses, we consider each step of the laboratory protocol (extractions, adapter ligations, and indexing) and assess the effect of host diet and taxonomy, extract pigmentation (suggestive of co-eluted secondary compounds), and sample input amount on protocol success rate. Finally, we assess if different types of samples differ with regards to the proportion of the metagenome that resembles oral microbiomes and the host DNA content. Our goal is to identify the most successful strategy for the study of dental calculus metagenomes across mammalian diversity and to help researchers and collection curators to optimise sampling and laboratory procedures for future dental calculus studies.

## Materials and Methods

### Sample collection and preparation for sequencing

Our dataset consists of 515 dental calculus samples belonging to 29 wild mammal species and three domesticates (Table 1), as well as 76 negative controls (50 from DNA extraction and 26 from library preparation, Supplementary Table 1). The samples were collected from natural history specimens housed in the following institutions: the Natural History Museum Vienna (Austria), the Royal Museum for Central Africa (Belgium), the Natural History Museum London (United Kingdom), National Museums Scotland (United Kingdom) and the Royal Belgian Institute of Natural Sciences (Belgium). Calculus was collected in a way that minimised contamination during the sampling process, using disposable lab coats, face masks and double gloves, with the upper layer of gloves changed before each new specimen. The bench surface on which sampling was carried out was decontaminated with bleach (at 10%) and rinsed with filtered water. Sampling was performed above a sheet of aluminium foil and calculus was collected onto Whatman Weighing Paper and transferred into 2 ml sterile tubes for storage at room temperature. Dental calculus chunks were dislodged from the teeth by applying pressure at the base of the visible deposit or, if present as a uniform film, gently scraped from the tooth surface using a sterile scalpel blade (Figure 1A), which was disposed of between samples. In rare cases, if the amount of material obtained was considered too low, samples from multiple individuals were collected in the same tube, matching individual provenance, sex and age as much as possible (Supplementary Table 1, column “Combined sample”).

**Figure 1.**
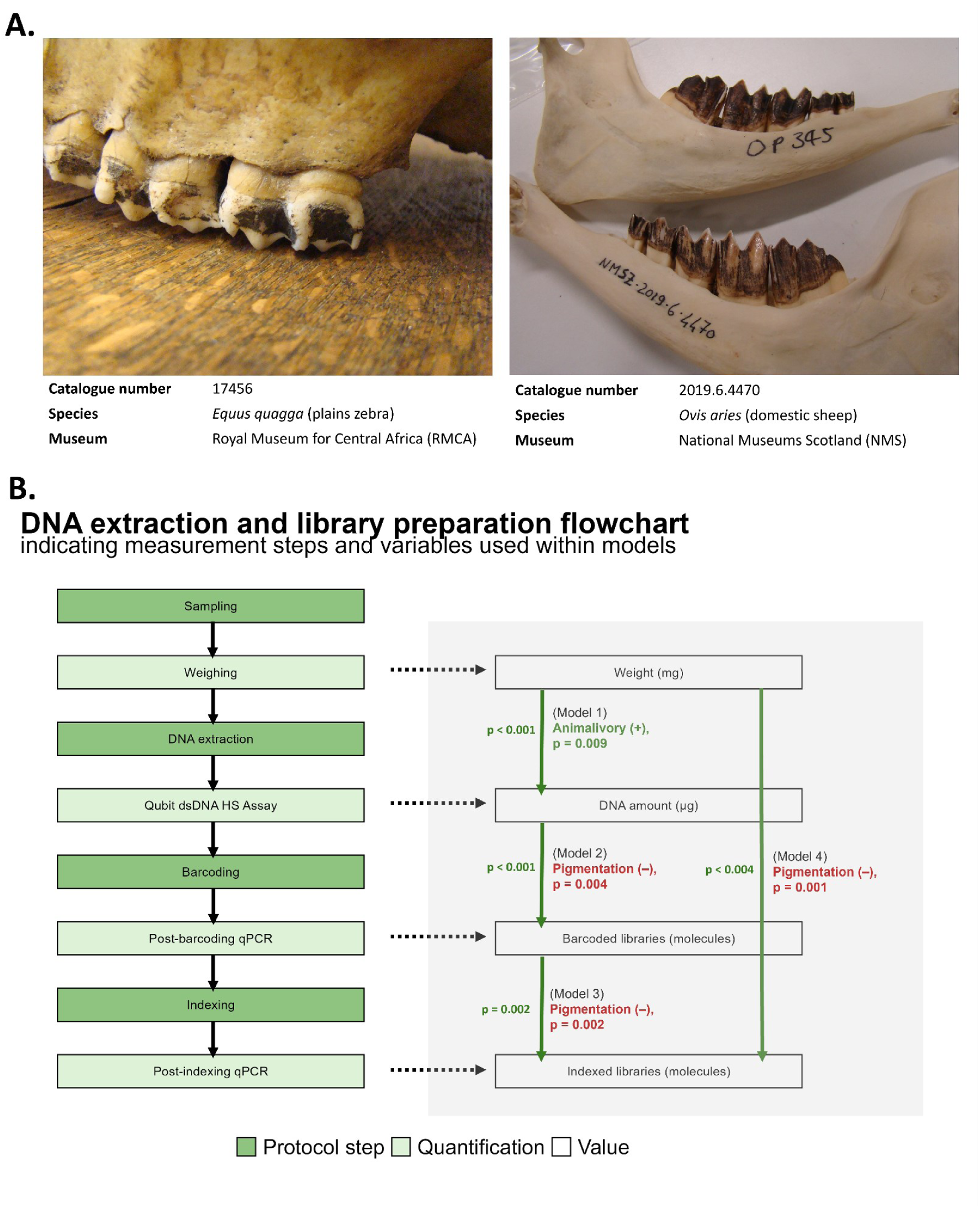
Dental calculus sampling, laboratory and statistical modelling procedures. **A.** Left: An example of ‘chunky’ dental calculus on a plains zebra, *Equus quagga*, specimen housed in the Royal Museum for Central Africa (RMCA). Right: an example of film-like dental calculus on a domestic sheep, *Ovis aries*, in National Museums Scotland (NMS). **B.** Left: Flowchart representing the laboratory protocol, including DNA extraction and Illumina sequencing library preparation. Dark green indicates protocol steps, whereas light green indicates the quantifications steps. Right: Variables used for the statistical models and tests of this study. Vertical arrows point from the variable used as a predictor (input) to the variable used as a response (output), with p-values signifying positive effects to the left of the arrow. Labels indicate any additional factors that were found to have a significantly positive effect (green) or significantly negative effect (red), alongside the corresponding p-values.

**Table 1.** List of species analysed in this study, alongside their taxonomic and dietary categories, and the number of extracted, barcoded, and indexed samples. Scientific names are according to (Mammal Diversity Database 2024), with the exception of the Malayan tapir, for which we use the updated name *Acrocodia indica* (Groves and Grubb 2011).

| Common name | Scientific name | Taxonomic order | Diet category | Samples extracted | Samples barcoded | Samples indexed |
| --- | --- | --- | --- | --- | --- | --- |
| African buffalo | *Syncerus caffer*<br>(Sparrman, 1779) | Artiodactyla | Herbivore | 10 | 10 | 8 |
| African elephant | *Loxodonta africana*<br>(Blumenbach, 1797) | Proboscidea | Herbivore | 15 | 13 | 13 |
| Alpine marmot | *Marmota marmota*<br>(Linnaeus, 1758) | Rodentia | Herbivore | 13 | 13 | 13 |
| Arabian camel | *Camelus dromedarius*<br>Linnaeus, 1758 | Artiodactyla | Herbivore | 9 | 9 | 9 |
| Argali | *Ovis ammon*<br>(Linnaeus, 1758) | Artiodactyla | Herbivore | 11 | 11 | 11 |
| Black rhinoceros | *Diceros bicornis*<br>(Linnaeus, 1758) | Perissodactyla | Herbivore | 2 | 2 | 2 |
| Bornean orangutan | *Pongo pygmaeus*<br>(Linnaeus, 1760) | Primates | Frugivore | 6 | 6 | 4 |
| Central African red colobus | *Piliocolobus foai* (de Pousargues, 1899) | Primates | Frugivore | 26 | 26 | 26 |
| Chamois | *Rupicapra rupicapra*<br>(Linnaeus, 1758) | Artiodactyla | Herbivore | 27 | 27 | 24 |
| Domestic horse | *Equus caballus*<br>Linnaeus, 1758 | Perissodactyla | Herbivore | 6 | 6 | 6 |
| Domestic pig | *Sus domesticus*<br>Erxleben, 1777 | Artiodactyla | Frugivore | 13 | 13 | 13 |
| Domestic sheep | *Ovis aries* Linnaeus, 1758 | Artiodactyla | Herbivore | 46 | 45 | 42 |
| Dugong | *Dugong dugon* (P. L. S. Müller, 1776) | Sirenia | Frugivore | 5 | 5 | 5 |
| Eastern gorilla | *Gorilla beringei*<br>Matschie, 1903 | Primates | Herbivore | 9 | 9 | 9 |
| Eastern grey kangaroo | *Macropus giganteus*<br>G. K. Shaw, 1790 | Diprotodontia | Herbivore | 14 | 14 | 14 |
| European badger | *Meles meles*<br>(Linnaeus, 1758) | Carnivora | Animalivore | 39 | 39 | 39 |
| European wild pig | *Sus scrofa* Linnaeus, 1758 | Artiodactyla | Frugivore | 26 | 26 | 26 |
| Giraffe | *Giraffa camelopardalis*<br>(Linnaeus, 1758) | Artiodactyla | Frugivore | 15 | 15 | 14 |
| Hippopotamus | *Hippopotamus amphibius* Linnaeus, 1758 | Artiodactyla | Herbivore | 13 | 11 | 11 |
| Indian muntjac | *Muntiacus muntjak*<br>(E. A. W. von Zimmermann, 1780) | Artiodactyla | Frugivore | 13 | 13 | 12 |
| Lowland paca | *Cuniculus paca*<br>(Linnaeus, 1766) | Rodentia | Frugivore | 13 | 12 | 12 |
| Malayan tapir | *Acrocodia indica* (A. G. Desmarest, 1819) | Perissodactyla | Frugivore | 8 | 8 | 7 |
| Muskox | *Ovibos moschatus* (E. A. W. von Zimmermann, 1780) | Artiodactyla | Herbivore | 6 | 4 | 0 |
| Okapi | *Okapia johnstoni* (P. L. Sclater, 1901) | Artiodactyla | Herbivore | 14 | 14 | 12 |
| Olive baboon | *Papio anubis*<br>(Lesson, 1827) | Primates | Frugivore | 21 | 21 | 21 |
| Orca | *Orcinus orca*<br>(Linnaeus, 1758) | Artiodactyla | Animalivore | 4 | 4 | 4 |
| Plains zebra | *Equus quagga* P.<br>Boddaert, 1785 | Perissodactyla | Herbivore | 27 | 27 | 27 |
| Roe deer | *Capreolus capreolus*<br>(Linnaeus, 1758) | Artiodactyla | Herbivore | 49 | 49 | 47 |
| South<br>American fur<br>seal | *Arctocephalus*<br>*australis* (E. A. W.<br>von Zimmermann,<br>1783) | Carnivora | Animalivore | 1 | 1 | 0 |
| South<br>American sea<br>lion | *Otaria flavescens* (G.<br>K. Shaw, 1800) | Carnivora | Animalivore | 16 | 14 | 14 |
| White<br>rhinoceros | *Ceratotherium*<br>*simum* (Burchell,<br>1817) | Perissodactyla | Herbivore | 13 | 13 | 13 |
| Yellow-backed<br>duiker | *Cephalophus*<br>*silvicultor* (Afzelius,<br>1815) | Artiodactyla | Frugivore | 25 | 25 | 25 |
| **TOTAL** |  |  |  | **515** | **505** | **483** |

The samples were processed in a specialised ancient DNA laboratory with strict anti-contamination protocols, following the procedure summarised in Figure 1B. DNA was extracted with a silica-based method developed for ancient DNA (Dabney et al. 2013) and slightly adapted for dental calculus (Brealey et al. 2020). In summary, dental calculus (with sample weight ranging from 1 to 35 mg) first underwent surface decontamination with a 10-minute UV irradiation followed by a brief wash with 0.5 M EDTA. The samples were then lysed overnight in a buffer consisting of 0.45 M EDTA and 0.025 mg/ml Proteinase K. The next day the lysate was mixed in 10 ml of binding buffer (5 M GuHCl + 40% (v/v) isopropanol + 0.05% Tween-20) and 400 μl of 3 M sodium acetate, and the DNA was bound to the silica membrane by centrifugation and washed twice using 450 μl of PE buffer (Qiagen) before being eluted with 45 μl of elution buffer consisting of EB (Qiagen) + 0.05% Tween-20. Samples were extracted in batches of up to 22 and two negative controls were processed in parallel for each sample batch, one at the start and one at the end of the batch. After elution, the extracts were quantified fluorometrically by a Qubit dsDNA HS Assay (Invitrogen).

Library preparation followed a double-barcoding double-indexing protocol to mitigate the effects of index-hopping (van der Valk et al. 2019). The DNA extract input was 20 μl. However, for some samples that showed inhibition, DNA extracts were diluted with ultrapure H_2_O (see Supplementary Table 1 for DNA input amounts). First, blunt-ended DNA fragments were created by T4 polymerase. Then, a barcoded adapter was added on each side of the fragment by T4 ligase and the gaps were filled in by Bst polymerase. We refer to this step as barcoding, otherwise known as adapter ligation (Meyer and Kircher 2010). The resulting barcoded libraries were quantified with qPCR and those with at least 10^6^ barcoded molecules underwent an indexing PCR to create complete Illumina libraries, so that each library molecule contained two barcodes and two indices (van der Valk et al. 2019). PCR cycles ranged from 10-14, depending on the qPCR results from the previous step, aiming to achieve similar final concentrations for all samples (Supplementary Table 1). A second qPCR was run to determine the number of fully-indexed library molecules, which were then pooled, aiming for equimolar amounts where possible. Sequencing was performed on four lanes of the Illumina NovaSeq X Plus platform (PE 150bp), aiming for ca. 10 million reads per sample.

### Dietary information

Since discrete categories such as carnivore, omnivore, etc. are insufficient in describing the dietary diversity among the included mammalian species, we used quantitative estimates for the percentages of different nutrients in the study species’ diet, as calculated by Lintulaakso et al. (2023) (Supplementary Table 2). These included crude protein, crude fibre, ether extract (proxy for fat content), nitrogen-free extract (proxy for non-fibrous carbohydrates, such as sugars and starch), and ash (proxy for inorganic compounds). The Lintulaakso et al. (2023) dietary database included all of the study species except for the domestic pig, *Sus domesticus*, so we used the same dietary data as for the European wild pig, *Sus scrofa*. We used the “calculated species main diet” from the Lintulaakso et al. database when discrete dietary categories were needed (Supplementary Table 2).

As some nutrients are strongly correlated, (e.g., protein and fat are both higher in an animalivorous diet, see Supplementary Figure 1), we performed a Principal Component Analysis (PCA) (Figure 2A) on the estimated dietary content of the study species, and extracted the first two principal components, which we scaled and centred on 0 to use as dietary predictors (Supplementary Table 2). Note that here we use animalivorous as a synonym for carnivorous to distinguish the dietary category from the taxonomic order Carnivora.

**Figure 2.**
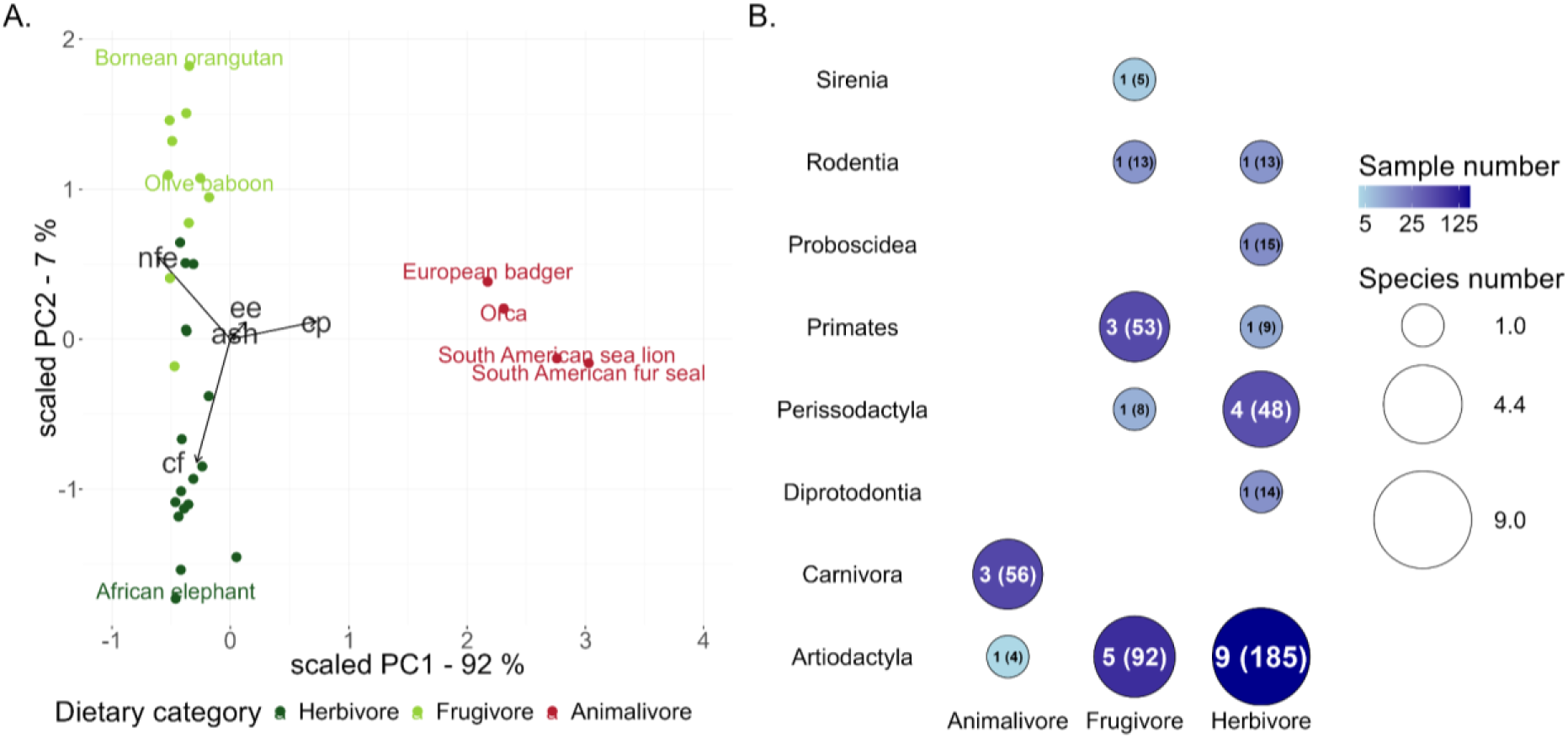
Summary of the dietary and taxonomic diversity in our dataset. **A)** Principal component analysis using proximate dietary data for the study species from Lintulaakso et al. (2023) (Supplementary Table 2), visualised using scaled principal components. The colours represent the calculated species main diet and the arrows indicate increase in the proportion of specific dietary components: ash = inorganic compounds, cf = crude fibre, cp = crude protein, ee = ether extract (proxy for fat), nfe = nitrogen-free extract (proxy for sugars and starches). Species labels are shown for all animalivores and also highlight the olive baboon, *Papio anubis* (which is assigned as “undetermined” in Lintulaakso et al., but was reassigned to frugivore because of its clustering with other frugivores), the African elephant, *Loxodonta africana* (as the species exhibiting the most herbivorous diet along the PC2 axis), and the Bornean orangutan, *Pongo pygmaeus* (as the species exhibiting the most frugivorous diet along the PC2 axis). **B)** Summary of the dataset (32 mammalian species, 515 samples), showing the number of species (circle size, number outside brackets) and number of samples (circle colour, number in brackets) for each dietary category and taxonomic order.

### Sequence pre-processing and source tracking

The sequences were demultiplexed based on their barcode-index combinations (Brealey et al. 2020), before trimming barcodes and adapters, removing low quality and short (<30 bp) reads, and merging forward and reverse reads using fastp (v. 0.23.4; Chen et al. 2018). We then mapped the metagenomes onto a combined reference of the human genome (GCA_000001405.29), PhiX (GCA_000819615.1), and the host species (Supplementary Table 3) using BWA-aln (v. 0.7.17; Li and Durbin 2009). Mapped reads were removed using samtools (v. 1.20; Danecek et al. 2021) and unmapped reads, which should be enriched for microbiome-derived sequences, were retained for downstream analysis. When a host reference genome was not available, we used a related species’ genome instead (Supplementary Table 3). To assess whether the proportion of host sequences preserved within dental calculus differs across taxonomically diverse hosts, we used a logistic regression with a quasibinomial distribution, and host order and diet as predictors. We evaluated the effects of these two factors using a Type II analysis of deviance on the logistic regression model, and performed pairwise comparisons across the levels of any significant factors, using generalised linear hypothesis testing implemented in the multcomp R package (v. 1.4.25; Hothorn, Bretz, and Westfall 2008).

To verify that our data indeed correspond to oral microbiomes, we used decOM (v. 1.0.0; Duitama González et al. 2023), a kmer-based microbial source tracking tool developed specifically for dental calculus metagenomes. We used the default reference to partition our metagenomes into the following microbiome sources; modern oral, ancient oral, skin, and sediment/soil (Supplementary Table 3). To identify factors that affect source proportions, we excluded species with less than four samples, recalculated the proportions, combined the ancient and modern oral proportions into a single value, excluded the “Unknown” partition and then performed a permutational analysis of variance (PERMANOVA), using the ‘adonis2’ function from the vegan package (v. 2.6.6.1; Oksanen et al. 2022) to assess the marginal effects of order and diet. We also specifically assessed if the oral microbiome proportion differs across host order and diet using the same approach as described above for the proportion of host DNA.

### Evaluating molecular protocol performance step by step

#### Linear models and statistical tests

The output of each step of the protocol (Figure 1B) was modelled using a weighted mixed-effects model implemented in the lme4 R package (v. 1.1.35.3; Bates et al. 2015) according to the general formula:

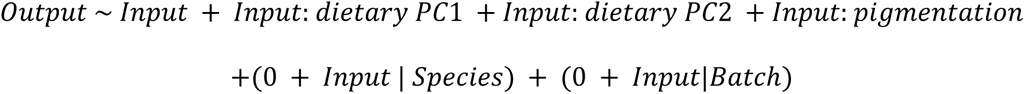

where *Output* and *Input* are continuous variables that differ for each processing step (i.e. for DNA extraction input was the sample weight in mg and output was the DNA amount in μg; see Figure 1B). The predictors *dietary PC*1 and *dietary PC*2 are the scaled principal components calculated based on the estimated dietary composition of the study species (Figure 2A), with PC1 primarily reflecting the amount of crude protein in the species diet and PC2 distinguishing between crude fibre and sugar/starch. Finally, *pigmentation* is a logical variable (TRUE/FALSE) indicating visible co-eluted components in the DNA extract, as detected by eye. *Species* and *Batch* indicate the host species and processing batch of the sample, respectively, and were included as random effects on the slope estimate. Specifically, *Batch* was included as a random effect to account for differences that could be due to technical factors in the laboratory, e.g., variation in buffer concentrations or incubation times. We used weighted least squares because our data were heteroscedastic (showing heterogeneous variance across the range of the response variable), to attribute less weight to observations with high variance.

As the indexing step of library preparation includes a PCR step, we do not expect a linear relationship between the input (expressed as the number of barcoded library molecules) and output (expressed as the number of indexed library molecules), because the PCR cycles vary between samples. Therefore, we calculated the PCR efficiency *E* of each reaction based on the following formula (Lalam 2006):

*N* = *N*_0_ × (1 + *E*)*^C^*, therefore 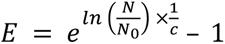, where *N*_0_ is the initial number of barcoded libraries, *N* is the final number of barcoded libraries and *c* is the number of indexing PCR cycles. We then used the same formula with the average *E* of all reactions to estimate the expected output of the indexing PCR, and we modelled the observed output as a function of the expected output.

To specifically test for the effect of different dietary categories (herbivore, frugivore and animalivore), we calculated the output to input ratio in each sample (e.g., for DNA extraction, the standardised output reflected the amount of DNA per μg of sample). We then tested how this standardised output differs across dietary categories using Kruskal-Wallis tests. If the global test was significant, we followed up with pairwise Wilcoxon tests, adjusting the p-values using the Holm method.

#### Machine learning

As library preparation is a costly and time-consuming procedure, we wanted to understand if specific sample characteristics are predictive of the probability that a given sample will produce usable Illumina libraries early in the sample processing procedure. Such relationships may depend on nonlinear interactions. For instance, there is, likely, an optimal DNA concentration for library preparation, with efficiency declining at both very low and very high input amounts, and this optimum may differ between different sample types. To identify such patterns, we performed binary classification in an attempt to predict the probability of success of the indexing protocol, using a machine learning approach, implemented in R using the mlr3 package (v. 0.20.2; Lang et al. 2019). We defined indexing failure when fewer than 10^9^ indexed molecules were produced, which corresponds to approximately the first quartile of indexed fragments for all negative controls (1.2 × 10^9^; Supplementary Figure 2A, Supplementary Table 1). As predictors we used DNA input (in μg), barcoded library volume used (μl), presence of extract pigmentation, host order and diet (expressed in scaled dietary PC1 and PC2; Figure 2). We first split our samples into a training (N = 324) and a testing dataset (N = 159), aiming for similar distributions of species in both sets (Supplementary Table 1). We tested learning algorithms (‘learners’) based on different approaches: k-nearest neighbours (‘kknn’), single layer neural network (‘nnet’), random forest (‘ranger’) and classification tree (‘rpart’) (Lang et al. 2024). We first optimised (‘tuned’) these learners by testing various hyperparameters using a Bayesian optimisation tuner (Schneider et al. 2024, 2) and evaluating them using repeated cross-validation (10 folds, 10 repeats) within the training dataset. After we identified the hyperparameter combinations that, for each learner, achieved the best predictions without overfitting, we evaluated the predictions of these tuned learners on the test dataset, using the log loss metric. A featureless learner, which always predicts the majority class from the data it is trained on (in our case, this was indexing success), was used to inform the minimum acceptable performance for the other learners. More details on the hyperparameters tested can be found in Supplementary Table 4. We finally used the best performing model, generated with a ‘ranger’ algorithm (an implementation of the random forest method) to estimate the relative importance of the predictor variables. We also generated predictions for indexing success using hypothetical samples – a combination of diet and range of sample weight that was not always present in our data. These consisted of DNA input amounts ranging from 0 to 0.5 μg, using brackets of 0.05 μg, for samples representing hypothetical members of the order Carnivora with animalivorous diets (scaled PC1 = 2.5, scaled PC2 = 0), Artiodactyla with herbivorous diets (scaled PC1 = −0.5, scaled PC2 = −1) and frugivorous primates (PC1 = −0.5, PC2 = 1).

## Results

### Dataset characteristics

Our dataset consists of 515 individuals from 32 mammalian species (Supplementary Table 1) with 1-49 samples per species (median = 13), representing eight taxonomic orders and a diversity of diets (Figure 2B, Table 1). As expected, most taxonomic orders have representatives from only one or two dietary categories, but our sampling was designed to limit intercorrelation of the two factors as much as possible.

A PCA based on the dietary composition of the study species (Supplementary Table 2) reveals two main clusters along the first principal component (PC1, Figure 2A). One cluster was characterised by high protein content and contained the European badger (*Meles meles*) and three marine animalivores. The second cluster contained herbivores and frugivores. The second principal component (PC2) distinguished herbivores with high fibre diets from frugivores with higher carbohydrate diets. Based on these groupings, we have amended the dietary classification for the olive baboon (*Papio anubis*) in our dataset, which was reported as “undetermined” in Lintulaakso et al. (2023), to “frugivore”.

We observed the presence of pigmented DNA extracts (usually with a brown or yellow hue) during sample processing, indicating the presence of secondary compounds co-eluting with DNA. Pigmented extracts were more common among herbivores (29.9%) and frugivores (19.9%) than in animalivores (1.6%) (Supplementary Figure 2B). The species with the largest proportion of pigmented extracts were the plains zebra (*Equus quagga*; 19 out of 27 samples), the muskox (*Ovibos moschatus*; 4 out of 6), the Indian muntjac (*Muntiacus muntjak*; 7 out of 13), the giraffe (*Giraffa camelopardalis*; 8 out of 15), the okapi (*Okapia johnstoni*; 7 out of 14), and the Malayan tapir (*Acrocodia indica*; 4 out of 8). Secondary compounds that produce pigmented DNA extracts can often act as inhibitors for enzymatic reactions such as PCR, therefore this factor was included as a predictor in the models.

### Host diet affects DNA concentration of dental calculus extracts

We explored the relationship between the weight of the sample used for DNA extraction and the resulting amount of DNA in the eluate, while accounting for the potential influence of host diet and extract pigmentation (included as interaction terms with weight) using a mixed linear model with weighted least squares (Model 1), after excluding one outlier sample with DNA output >4 μg.

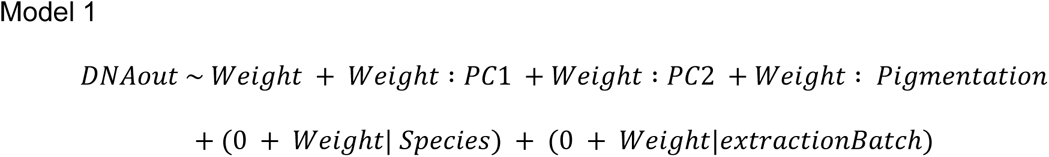

where *DNAout* refers to the extracted DNA amount (in μg) and *Weight* refers to sample weight used as input (in mg). As our model did not fit the assumptions of normality of variance (Shapiro-Wilk normality test: W=0.85, p < 0.001, Supplementary Figure 3), we used weighted least squares instead of ordinary least squares. As expected, we found that sample weight had a positive effect on extracted DNA amount (p < 0.001), suggesting that for every mg of dental calculus sample we extract approximately 0.023 μg of DNA (Figure 3A, Supplementary Table 5). We also found a positive interaction with dietary PC1 (which reflects the carnivory levels) with an interaction coefficient of 0.009 (p = 0.01). This suggests that we retrieve more DNA per mg of sample for more carnivorous taxa. According to our estimates, we can expect 0.046 μg of DNA per mg of dental calculus for a typical carnivore (with PC1=2.5, see Figure 3A), but only 0.018 μg per mg of sample for a herbivore (PC1=−0.5) (Figure 3A). The remaining predictors did not have a significant effect (Supplementary Table 5).

**Figure 3.**
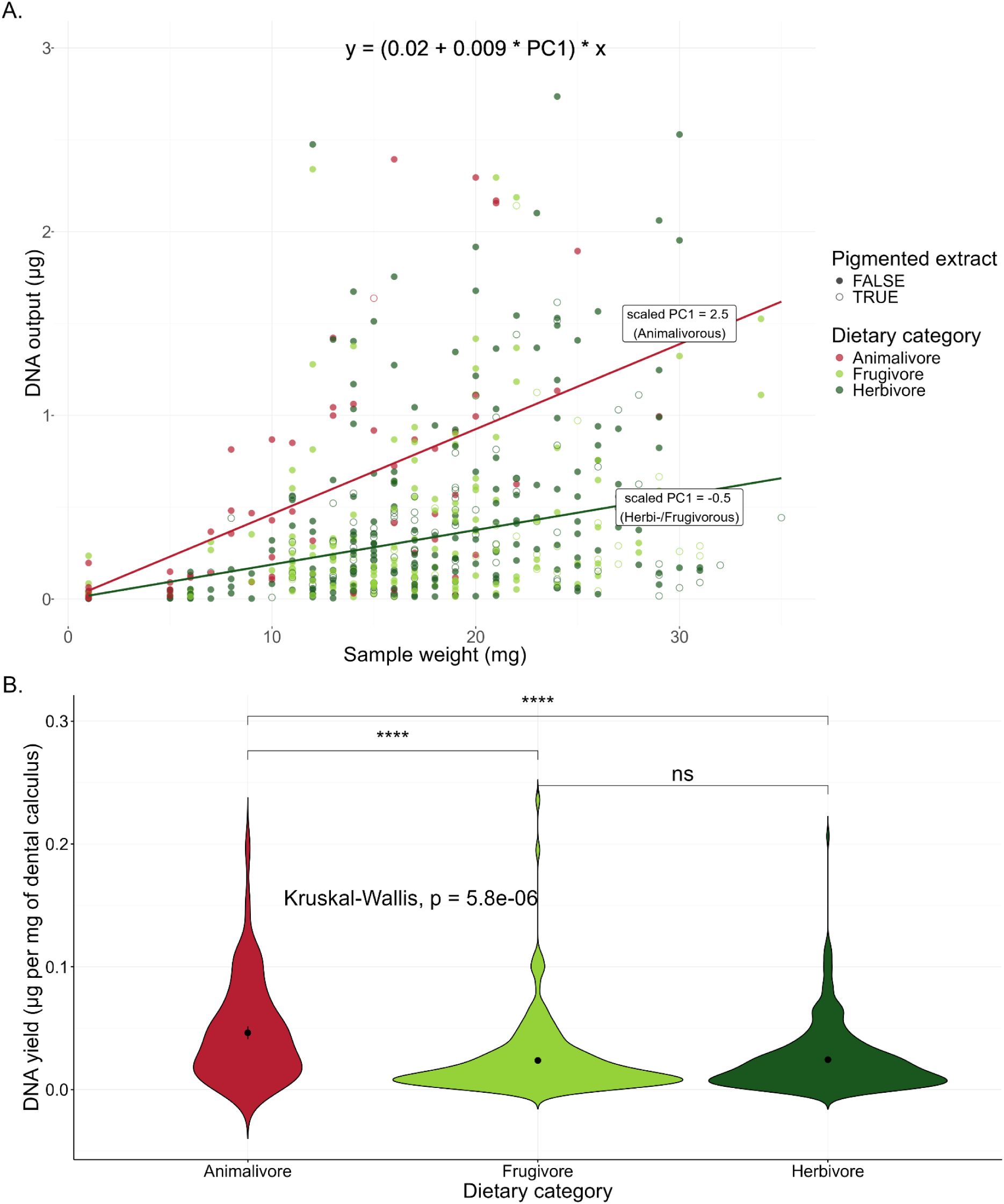
Factors affecting DNA output following dental calculus DNA extraction. **A)** The relationship between sample weight (in mg) and extracted DNA (in μg) for 515 samples. The formula on the top is based on all significant model estimators and the two lines show the expected DNA concentration for a species with scaled PC1 = 2.5 (indicative of a carnivorous diet) and a species with a scale PC2 = −0.5 (non-carnivorous diet). **B)** DNA yield for different dietary categories. Significance levels:(****) p < 0.0001, (np) p > 0.05.

We also tested for differences in DNA yield (expressed as proportion of sample weight) between discrete dietary categories (Figure 3B). Indeed, diet category had a significant effect on the DNA yield (Kruskal-Wallis test: p < 0.001), with animalivore hosts having more DNA per mg of extracted dental calculus, confirming the above results.

### Pigmented dental calculus extracts negatively affect library preparation

The 505 samples that were retained after DNA extraction underwent barcoding, during which a double-sided barcode was added to each DNA fragment. We modelled this first library preparation step (barcoding) as a linear mixed model with weighted least squares (to account for heteroscedasticity; Supplementary Figure 4), similar to the formula used to model DNA extraction. Specifically for the barcoding step, we first excluded seven outliers with barcoding outputs larger than 8×10^10^ copies and one with a DNA input amount larger than 3μg, and used the following formula:

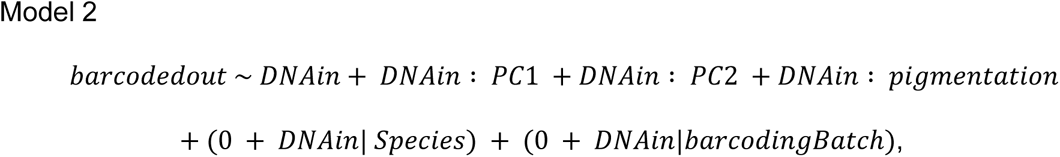

where *barcodedout* is the output of barcoded libraries expressed in 10^10^ molecules and *DNAin* is the input DNA in μg. As expected, we found a positive relationship between DNA input and barcoding output with a slope of 2.8 (p-value < 0.001), but this relationship was affected by extract pigmentation with an interaction of −2.2 (p-value = 0.004) (Supplementary Table 5), suggesting that, when dealing with pigmented extracts, increasing the DNA input amount may only have marginal or no benefits for the barcoding output (Supplementary Figure 4A, Supplementary Table 5). We also found a marginal effect of the dietary PC2 (frugivory versus herbivory axis) suggesting that herbivores may be generating a lower barcoding output than frugivores (Supplementary Table 5). This became more evident when discrete dietary categories were compared. Clear DNA extracts from herbivores have a lower barcoding output than extracts from hosts with other diets (pairwise Wilcoxon tests: p < 0.01, Figure 4A).

**Figure 4.**
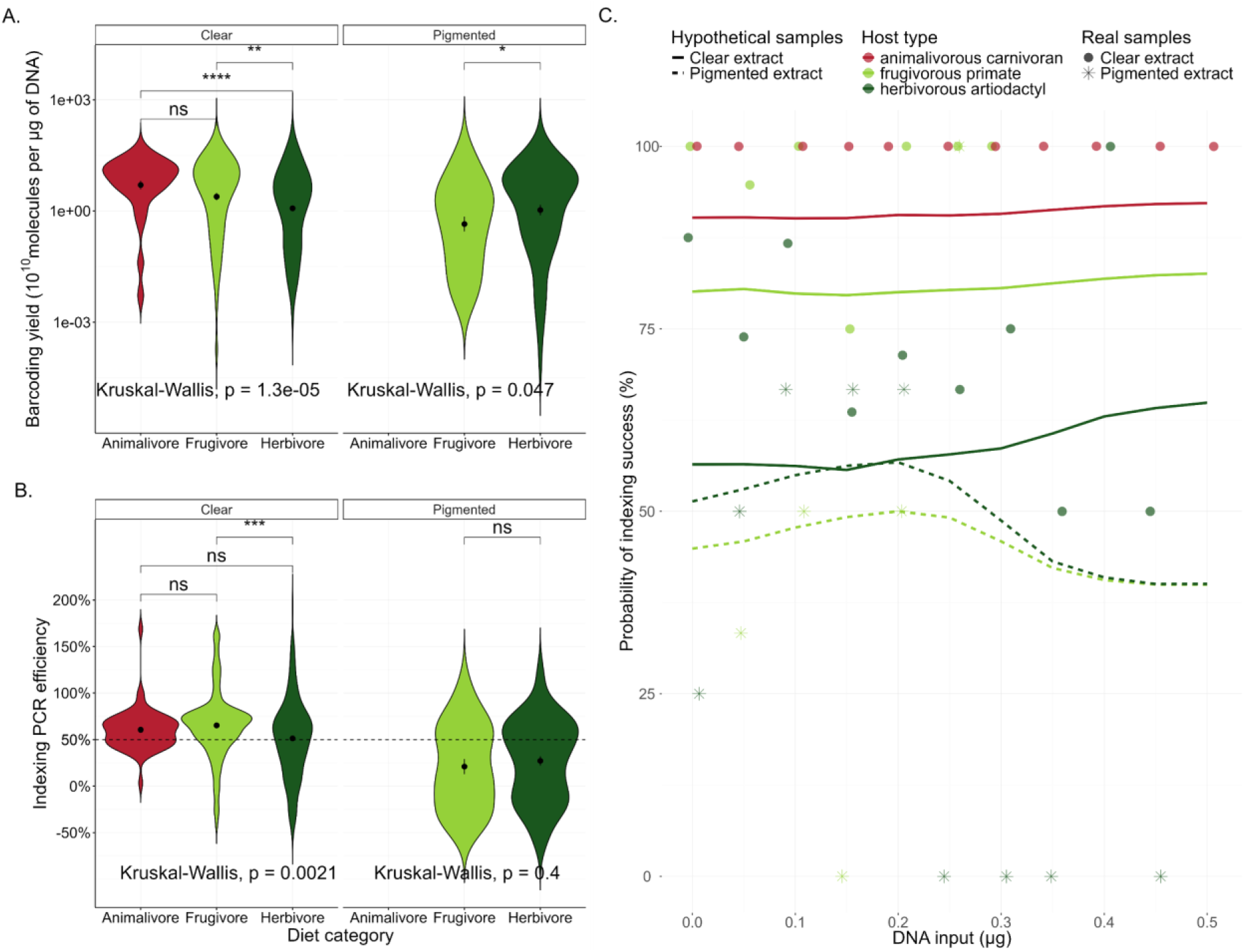
Outcome of library preparation by dietary category and extract pigmentation. **A)** The amount of barcoded libraries (in 10^10^ molecules) per μg of input DNA. Note the logarithmic scale of the y-axis. **B)** The estimated indexing PCR efficiency per sample. Under ideal conditions the efficiency would be 100%, indicating a doubling of DNA fragments at each cycle. The horizontal dashed line indicates the average efficiency (49.9%) across all indexed samples in our dataset. The effect of diet category was assessed using Kruskal-Wallis tests separately for clear and pigmented extracts, pairwise comparisons were performed with Wilcoxon tests and p-values were adjusted using the Holm method. Significance levels:(****) p < 0.0001, (***) p < 0.001, (**) p < 0.01, (*) p < 0.05, (ns) p > 0.05. **C)** Indexing success probability (displayed as lines) as predicted by a ranger classifier trained on a subset of our data. The predictions were generated based on three hypothetical host species: animalivorous carnivorans (scaled PC1 = 2.5, scaled PC2 = 0, red lines), herbivorous artiodactyls (scaled PC1 = −0.5, scaled PC2 = −1, dark green lines) and frugivorous primates (scaled PC1 = 0.5, scaled PC2 = 1, light green lines). We assumed DNA input amounts spanning every 0.05 μg from <0.01 to 0.5 μg for both clear and coloured extracts. The coloured circles show observed indexing success values of similar hosts: animalivorous carnivorans represented by the badger and the South American sea lion (48 samples), herbivorous artiodactyls represented by the Arabian camel, the roe deer, the hippopotamus, the okapi, the argali, the domestic sheep, the chamois and the African buffalo (146 samples), and frugivorous primates represented by the olive baboon, the Central African red colobus and the orangutan (50 samples). The real indexing success was calculated within bins of DNA input μg that approximated the values used for the hypothetical data.

After barcoding, 22 libraries were excluded because of too low numbers of library molecules (less than 100x barcoded molecules compared to the negative control in that library batch). Of these 63.6% (n=14) had a pigmented extract, whereas only 21.8% (n=104) of the retained 475 barcoded libraries did, suggesting that extract pigmentation may reflect the presence of inhibitors, which impede the barcoding step of the protocols.

The retained libraries underwent an indexing PCR, with the number of PCR cycles adjusted according to the template concentration (Supplementary Table 1). We found that a considerable proportion of the dataset failed to amplify as expected, with 100 (20.7%) indexed libraries producing fewer than 10^9^ copies. For comparison, negative controls, which contain no sample and start at a much lower concentration, produced a median of 4 × 10^9^ indexed copies (range: 3 × 10^8^ - 1 × 10^12^) (Supplementary Table 1). Of these failed libraries (with < 10^9^ indexed copies), 47% (n=47) had a pigmented extract, whereas from the successful libraries (> 10^9^ indexed copies), only 15.1% (n=58) did (Supplementary Figure 2B).

We estimated the efficiency of the indexing PCR to be on average 49.9% (reflecting an increase of DNA molecules by 149.9% at every cycle). However, we found that pigmented extracts showed lower PCR efficiency (mean: 24.8%) compared to clear extracts (mean: 57.8%, Kruskal-Wallis test: p < 0.001, Figure 4B, Supplementary Figure 5B). Even amongst clear extracts herbivores tended to have somewhat reduced efficiency (Figure 4B), similar to what we observed at the barcoding step. To better understand the relationship between the observed (*indexout*) and the expected (*exp*) indexed molecules (under the assumption that all samples have an average PCR efficiency of 49.9%), we implemented a linear mixed effects model (Model 3) with weighted least squares (once again to account for heteroscedasticity in the data; Supplementary Figure 6), after removing three outliers with more than 400 × 10^10^ observed indexed molecules.

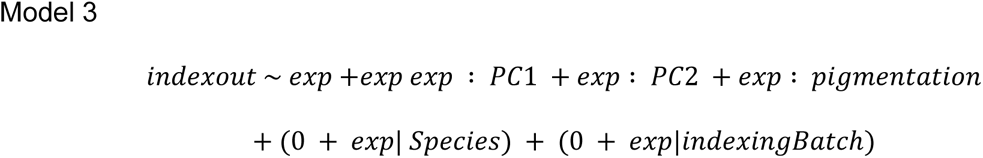

We observed, as expected given how we calculated the expected indexed molecules, a positive relationship between the observed and expected indexing with an effect size of 1.50 (p = 0.002), very close to the average PCR efficiency. However, we found a negative effect of pigmentation equal to −0.67 (p = 0.002, Supplementary Figure 5B, Supplementary Table 5). We did not detect an effect of diet as expressed by dietary principal components.

At the end of the entire protocol, 383 successfully indexed libraries were obtained from the initial 515 extracted samples, indicating an overall success rate of 74.4%. In an attempt to assess the likely success of the laboratory methods as early as possible during sample processing and derive general recommendations from the moment of sample collection, we ran a weighted mixed effects model to determine the impact of sample input weight on the final indexing output:

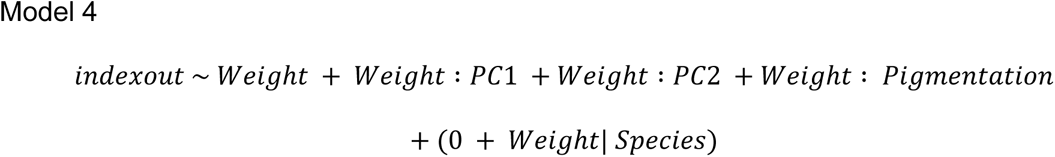

We found a small positive effect of sample weight, suggesting that the indexing output increases by 1.49 x 10^10^ indexed molecules for each μg of sample (the average output is 42.38 x 10^10^), as well as a negative effect of extract pigmentation (Supplementary Table 5).

Although we consistently observed that input had a positive impact on the output of each protocol step, it is possible that this relationship is nonlinear, as there may be an optimal amount of DNA input, which can vary depending on the presence of inhibitors. To investigate such nonlinear relationships, and in the interest of planning future research, we employed a machine-learning method to model the success probability of the indexing step, using information that is available at the DNA extraction stage. Specifically, given the high failure rate at the indexing step (N = 100, or 20.7% with less than 10^9^ molecules; Supplementary Figure 2A), we performed binary classification using as predictors the following variables: DNA input (μg), host order, host diet expressed quantitatively as scaled PC1 (carnivory axis) and PC2 (herbivory vs frugivory axis), presence/absence of extract pigmentation, and barcode library volume (μl). The minority class (indexing failure) was coded as TRUE and the majority class (indexing success) was coded as FALSE. The best performing learner was the ranger algorithm (with hyperparameters mtry = 3, num.trees = 985, min.node.size = 20, max.depth = 18; Supplementary Table 4). However, even this best-performing learner had a relatively small predictive power, with an Out-of-Bag (OOB) of 0.12, moderate false-negative rate (FNR = 0.02), but rather high false-positive rate (FPR = 0.85). The high FPR shows that the model predicts some successful samples to fail indexing and may be partly due to the relatively low number of true positives in our dataset (21% of samples). The resulting model suggests that the most important factors determining indexing output are, in descending order, extract pigmentation, dietary PC2 (herbivory vs. frugivory axis), dietary PC1 (the carnivory axis), barcoded library input, taxonomic order and finally the DNA input (Table 2).

**Table 2.** Variable importance for indexing success/failure, as estimated using a ‘ranger’ (random forest) machine learning algorithm.

| Variable | Importance |
| --- | --- |
| Extract pigmentation | 8.57 |
| Dietary PC2 | 6.58 |
| Dietary PC1 | 5.56 |
| Barcoded library volume (μl) | 5.14 |
| Host taxonomic order | 4.34 |
| DNA input (μg) | 3.15 |

To better understand how these predictors affect indexing success rate, we used the trained model to make predictions for hypothetical samples. We provided values representative of the three different dietary categories, with DNA input ranging from <0.01 to 0.5 μg and with clear and pigmented extracts (Supplementary Table 6). The ranger model agreed with the linear models (Models 2 and 3) in that extract pigmentation and herbivory adversely affect library preparation. Indexing success for pigmented and/or herbivorous dental calculus samples was predicted to be between 42.1% and 69.3%, whereas for clear extracts from animalivorous or frugivorous species it was between 78.7% and 94.5% (Figure 4C). Beyond the insights obtained from the linear models, the machine learning predictions suggested that for samples with pigmented extracts, increasing the input amount beyond approximately 0.2 μg of DNA reduces the predicted success rate (Figure 4C, Supplementary Table 6). This may reflect the inhibitor content increasing with higher sample input amounts. However, these predictions should be considered with caution, as the model error rate was high, and the observed success rates varied considerably (Figure 4C, data in Supplementary Table 6). In particular, for samples from herbivorous hosts with clear extracts, the real data suggest that indexing success decreases with DNA input, which is not reflected by the machine learning model.

### DNA input has low impact on library complexity

So far we have explored how the amount of DNA used for library preparation affects its efficiency and success rate. However, a concern in metagenomics, especially when working with degraded ancient or historical samples, is the information content of a sample (often referred to as library complexity), which is related to diversity of DNA fragments. In this sense, DNA input can be viewed as ‘sampling effort’, with sequencing libraries generated from low-input DNA showing lower complexity and hence fewer unique library molecules.

As a proxy for library complexity we investigated the percentage of unique reads after sequencing and how it is affected by DNA input. We considered a subset of 303 samples that achieved a sequencing depth between 10^6^ and 2×10^7^ reads. The percentage of unique reads reaches a plateau at around 95% once DNA input reaches 0.01 μg (Supplementary Figure 7). Sequencing depth can also impact the percentage of unique reads (even with low input, a very shallowly sequenced library will have mostly unique reads) and since our samples varied considerably in sequencing depth, we also inspected subsets of samples with more similar sequencing depths. We split our data into four subsets, each reflecting a quartile of sequencing depth (Supplementary Figure 8) and find roughly the same pattern for the proportion of unique reads as in the complete dataset. Overall, for DNA input > 0.01 µg, 80.8% of samples (227 out of 281 samples) showed >90% unique library molecules. For samples with DNA input < 0.01 µg, only 27.3% of samples (6 out of 22) reached this library complexity. The sequencing performed in this study is shallow by many metagenomics standards and therefore for more deeply sequenced data, higher DNA input may be necessary to maintain sufficiently complex libraries for downstream analysis. Nevertheless, this analysis demonstrates that the DNA input amount considered as ideal for successful library preparation (0.2 µg; Figure 4C) is sufficient to obtain sequencing libraries with good complexity. Considering the estimated average yield of 0.023 μg of DNA per mg of dental calculus, this DNA amount corresponds to approximately 8.69 mg of sample.

### Host and oral microbiome content vary across mammals

In total 401 indexed dental calculus libraries were sequenced on an Illumina NovaSeq platform. After pre-processing the metagenomic reads, each sample had on average 6.0 million reads (and up to 54.3 million, Supplementary Table 3). We mapped the metagenomic reads to the host and human genome and, after excluding the human-mapped sequences as contamination, we calculated the percentage of host reads, which varied from 0% to 83.63% with an average of 5.9% (Supplementary Figure 9). The analysis of deviance suggested a significant effect of host order (Type II analysis of deviance: chi-squared = 22.1, p-value = 0.002), but not host diet (chi-squared = 1.19, p-value = 0.55) in determining the proportion of host reads. Pairwise comparisons between host orders showed that African elephants (order Proboscidea) generally had higher host content than most other hosts, with the exception of kangaroos (order Diprotodontia) and rodents, which also had relatively high host content (Supplementary Table 7, Supplementary Figure 10). Sirenia had consistently the lowest host content (0.05-0.5%, median 0.1%; Supplementary Figures 9 and 10), but were not significantly different from other host orders, likely due to their small sample size (N = 5). Primates also tended to have fewer host reads (Supplementary Figures 9 and 10). These results may reflect technical artefacts rather than differences in the intrinsic properties of dental calculus (see Discussion).

After removing host and human reads, we estimated the oral microbiome proportion among the remaining sequencing reads. To this end, we used decOM for 27 species with at least five samples each (337 samples in total), confirming the presence of oral microbiome signature in all species (Supplementary Table 3, Supplementary Figure 11). On average, 43.9% of the metagenomes was estimated to resemble the oral microbiome (ancient or modern) and another 44.4% was of unknown origin (i.e. not matching any of the included sources, Figure 5). Skin and soil/sediment microbiota, which are likely contaminants (although soil could be co-ingested during feeding), accounted for an average of 5.5% and 6.0%, respectively (Supplementary Table 3). The high unknown proportion is expected in wild animals with uncharacterised microbiomes. The two species with the largest metagenomic proportion of unknown source were both marine mammals, the dugong (*Dugong dugon*: on average 83% unknown) and the South American sea lion (*Otaria flavescens*: 71.8% unknown, Figure 5).

**Figure 5.**
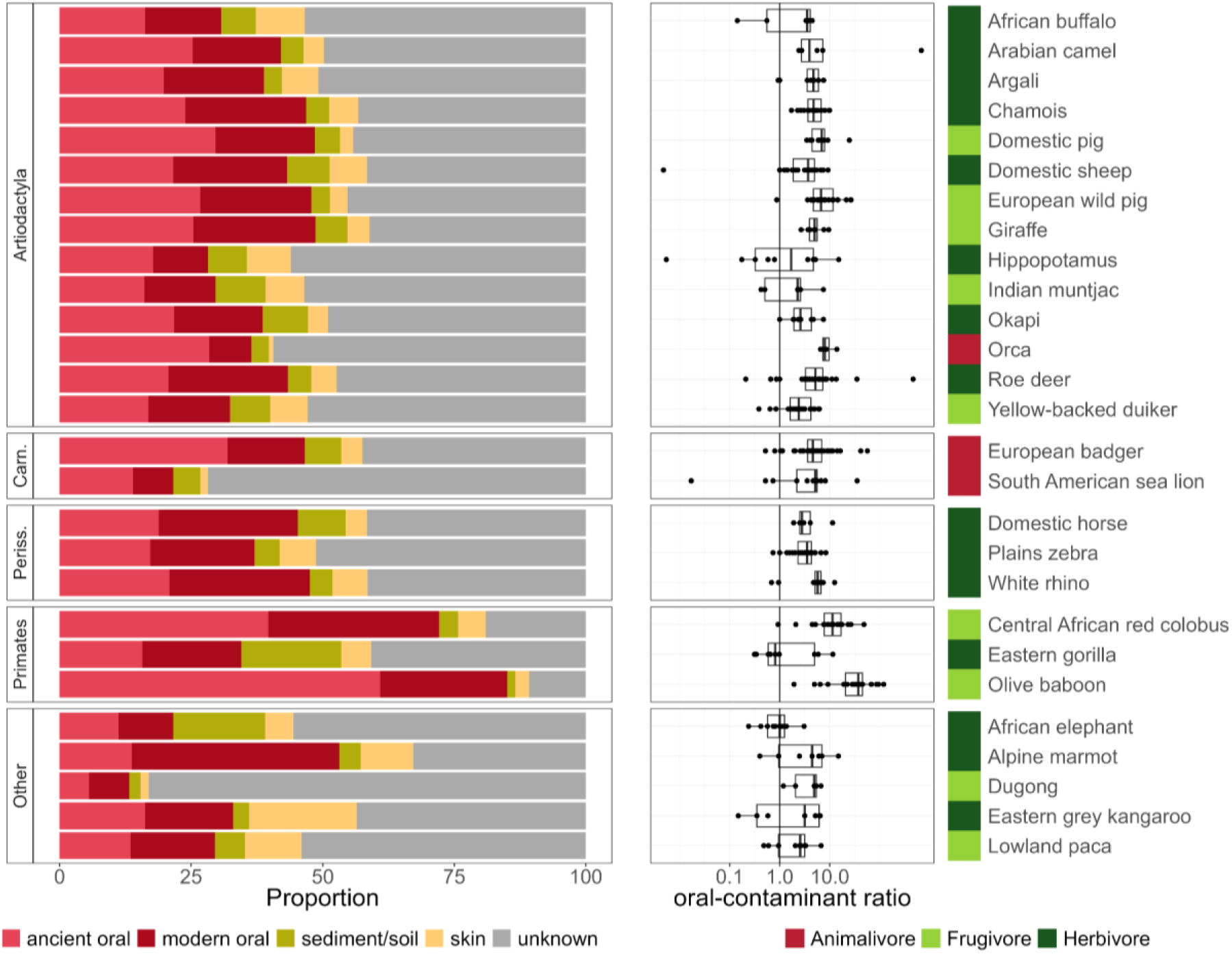
Source tracking of the dental calculus metagenomes. Left (barplot): Estimated source composition of dental calculus metagenomes (after removing reads mapping to the host and human genome) averaged across samples for each host species. Analysis based on 376 samples from 27 species with at least five samples each (see representation per sample in Supplementary Figure 11). The “Unknown” partition refers to kmers not represented in the included sources. Plot data can be found in Supplementary Table 3. Right (boxplot): Oral (ancient and modern) to contaminant (skin and soil) ratio per species. Vertical line indicates a ratio of 1. Note the logarithmic scale on the x-axis. The coloured bar at the right-hand side indicates mammalian host diet.

Disregarding the unknown fraction, we find that the composition of dental calculus microbiomes varied across orders (PERMANOVA marginal effects: R^2^ = 0.11, p-value = 0.001) and to a lesser extent by dietary category (R^2^ = 0.02, p-value = 0.01). Similarly, the oral microbiome proportion varied between taxonomic orders (Type II analysis of deviance: chi-squared = 567.0, p-value < 0.001) and to a lesser extent diets (chi-squared = 20.7, p-value < 0.001). Oral microbiome proportion was greater than the proportion of likely contaminants (skin and sediment/soil microbiota combined) in most samples and species (Figure 5). We found that Primates had a higher oral microbiome proportion compared to all other host orders (Generalised linear hypothesis testing: 2.9 ≤ ratio ≤ 16.5, p-value < 0.001) (Supplementary Table 8, Figure 5, Supplementary Figure 11). On the other hand, dugongs (*Dugong dugon*), the only representative of the order Sirenia, had the lowest oral microbiome proportion (0.061 ≤ ratio ≤ 0.45; p-value < 0.05), followed by the African elephant (*Loxodonta africana*), the only representative of the order Proboscidea (0.13 ≤ ratio ≤ 2.2, p-value < 0.05; the only ratio larger than 1 reflects the comparison against Sirenia) (Supplementary Table 8, Figure 5). It is noteworthy that the 15 African elephants in our dataset were collected at the same time and place (Supplementary Table 1), and were likely treated and housed identically. Therefore, these results could reflect the low preservation of these particular elephant specimens and not a species-specific characteristic. This is not the case for five dugong specimens from National Museums Scotland, whose collection dates span over a century.

## Discussion

Dental calculus metagenomics is an untapped resource for microbiome studies across a diversity of mammals (Brealey et al. 2020; Moraitou et al. 2022; Ottoni et al. 2019; Fellows Yates et al. 2021). With growing interest in this field, it is crucial to devise optimal processing methods. Until now metagenomic analyses of dental calculus have been carried out in only a few mammalian species, with a narrow representation of ecological (e.g., diet, lifestyle) and taxonomic diversity. In this study we massively extend this diversity, by analysing dental calculus from 32 species across the mammalian tree of life, including marine, aquatic and terrestrial taxa with representatives from eight extant mammalian orders. We recovered an oral microbiome signature from all studied species, confirming the utility of dental calculus for oral microbiome investigations in many previously unstudied species, but also found that not all host species perform equally well in the laboratory.

The most striking, albeit not unexpected result, is the severely detrimental effect of DNA extract pigmentation on the efficacy of the molecular laboratory protocols, particularly the library preparation steps. Pigmented extracts produced fewer barcoded and indexed molecules (Figure 4) and were removed from the experiment at a higher rate due to this lower performance. DNA extract coloration has been linked to the presence of secondary compounds that negatively impact the enzymatic steps of molecular laboratory workflow, particularly pectins, polyphenols, polysaccharides, and humic acid (Sidstedt et al. 2015; Loomis 1974; Sidstedt, Rådström, and Hedman 2020). Plants are particularly rich in inhibitors (Schrader et al. 2012; Demeke and Adams 1992; Winkel-Shirley 2001), which may explain why herbivores and frugivores showed a higher proportion of pigmented DNA extracts than animalivores (Supplementary Figure 2B). Tropical and subtropical plants are especially enriched in compounds, such as phenols, to defend themselves against herbivory (Becerra 2015) and, correspondingly, the species with the largest proportion of pigmented extracts in our dataset (plains zebra, muntjac, giraffe, okapi and malayan tapir, muskox) were almost exclusively tropical/subtropical browsers and/or grazers (with the exception of the muskox).Therefore, it seems that enzymatic inhibition in dental calculus samples may be caused by a diet rich in secondary compounds. Even without evident pigmentation, a herbivorous diet had a negative effect on the efficacy of the barcoding and indexing steps (Figure 4, and Supplementary Figure 5). As extract pigmentation was recorded by visual inspection, low concentrations of inhibitors may have remained undetected, but were likely present in herbivorous hosts.

The presence of inhibitors poses challenges not only in studies of dental calculus but also other historical and ancient samples, including sediments (Nota et al. 2022; Rayo et al. 2022). Sample dilution, which is often employed to ‘dilute out’ the inhibition, can be prohibitive for very degraded, low-quantity material. Our findings suggest that this may be a good practice for samples with a high inhibitor load: pigmented extracts have a higher predicted success rate for DNA inputs below 0.2 μg (Figure 4C, Supplementary Table 6). In addition, we retrieved >90% unique reads for most libraries with at least 0.01 μg of DNA input (Supplementary Figures 7 and 8). In conjunction, these findings suggest that diluting to a final concentration between 0.01 and 0.2 μg (10-200 ng) may help evade inhibition without sacrificing library complexity. Assuming the average DNA yield of 0.023 μg per mg of dental calculus (based on Model 1), this indicates that using more than 8-9 mg may increase the chance of enzymatic inhibition. However, it is important to remember that we performed relatively shallow sequencing, and that studies that require deeper sequencing will likely need a larger quantity of complex libraries. Furthermore, our samples were relatively young, mostly from the 19th and 20th centuries (Supplementary Table 1), and DNA from older specimens is likely to be lower in quantity and more degraded. Therefore, a better understanding of the inhibition mechanism is required to develop more robust methods for dental calculus metagenomics, e.g., by relying on inhibition-tolerant DNA polymerases (Sidstedt et al. 2015) or amplification facilitators (Wilson 1997).

In addition, we found that dental calculus from different species varied in the amount of DNA that can be extracted from it. Animalivores had a higher DNA yield relative to sample weight, 0.042 μg of DNA per mg of dental calculus, compared to 0.015 μg for a typical herbivore. This is a considerably lower yield than, e.g., modern dental pulp (~0.200 μg/mg), but at least for carnivorous taxa, it is comparable to that of blood (~0.030 μg/mg) (Siuta et al. 2024). This distinction may reflect intrinsic differences in DNA content of dental calculus depending on the dietary composition. As a living biofilm, dental plaque (the uncalcified form of dental calculus) consists of bacterial cells that are embedded within a secreted extracellular matrix (Schroeder and Shanley 1969). The composition of this biofilm, and specifically the matrix-to-cell ratio, can vary (Frank and Brendel 1966) and hence lead to variation in DNA content per gramme of material. Polysaccharides, including a fraction from diet, are an important component of the dental plaque matrix (Jakubovics et al. 2021; Frank and Brendel 1966). This may explain why carnivorans in particular, with lower carbohydrate dietary content (Supplementary Figure 1), have a more “densely packed” dental calculus with a larger number of cells, resulting in higher DNA yield. Alternatively, this observation could reflect inhibition during DNA fluorometry-based quantification, which may lead to underestimation of DNA concentration in frugivores and herbivores (Zipper et al. 2003; Sidstedt et al. 2015). As dental calculus tends to be rarer in wild animalivores than in herbivores and, if present, is found in lower quantities (Richards et al. in review), a higher DNA yield in animalivores is encouraging, as it suggests that less material can be used to obtain sufficient DNA for downstream applications.

Dental calculus is known to entrap host and dietary molecules. However, the initial excitement about the possibilities of studying host genomics and diet from this material has been subdued after the realisation that eukaryotic reads account for <1% of DNA obtained from human dental calculus (Mann et al. 2018). Although higher host content was detected in non-human mammals (from 0 up to 74%; Brealey et al. 2020; Moraitou et al. 2022), so far most host genomic analyses from dental calculus samples were primarily focusing on mitochondrial information (but see Brealey et al. 2020). Our results suggest variation in host content across mammalian orders, with a lower proportion in primates and a higher proportion in elephants, and to a lesser extent kangaroos and rodents (Supplementary Table 7, Supplementary Figures 9 and 10). However, these results should be interpreted with caution. There is a large variation in morphology of dental calculus (Richards et al. in prep), which impacts the ease of sampling. Chunky calculus deposits, such as frequently seen in omnivores and carnivores (and some herbivores, Figure 1) can be removed by applying pressure at the base of the deposit, whereas filmy deposits require scraping of the tooth surface, thus increasing the probability of co-sampling host tooth tissues. The sampled elephants tended to have less dental calculus (Richards et al. in review) than other taxa. Therefore, during sampling, more enamel material may have inadvertently been collected alongside the calculus. Similarly, due to the small size of rodents, more teeth had to be sampled, again increasing the possibility that enamel was collected. As a result, host tissues may have been co-extracted with calculus, thus leading to higher host read proportions. On the other hand, the lower host content in primates may be the result of a technical artefact during bioinformatic analysis. Specifically, to remove host reads from the dental calculus metagenomes, we performed competitive mapping, using a concatenated reference of the host and the human genome (the latter used as the most likely contaminant during museum storage, handling, and laboratory processing). However, in the case of primates the genetic similarity between human and primate host may lead to increased removal of primate reads that map to the human genome, particularly if the quality of the primate reference is low. Such instances will be less frequent for host species that are evolutionarily more distant from humans, e.g., carnivorans.

A similar explanation may also apply as to why primates show by far the highest oral proportion (Figure 5, Supplementary Figure 11). Microbial source tracking relies on known references, which in most cases, including those used here, are human microbiomes (Duitama González et al. 2023). Primates likely show greater similarity of oral microbiota to those of humans, which leads to a higher proportion characterised as oral instead of “unknown”. The two species with the highest “unknown” proportion in their microbiomes are both marine: the South American sea lion (*O. flavescens*) and the dugong (*Dugong dugon*). Oral microbiome studies of other marine mammals, such as bottlenose dolphins, *Tursiops truncatus*, reveal an abundance of uncharacterised taxa, not just belonging to unknown species or genera, but entirely novel phyla (Dudek et al. 2017). Therefore, it is expected that human oral microbiomes do not bear high similarity to those of marine mammals. This result is yet another reminder of the considerable gaps in omics databases that are so often used as reference and highlights the need to expand the characterisation of the animal microbiomes to a wider phylogenetic and ecological diversity and to develop and embrace methods that do not rely solely on reference databases (Worsley et al. 2024; Leonard et al. 2024).

Our results highlight several key considerations relating to dental calculus samples that we hope will be useful for researchers and curators alike when planning future projects. First, despite the considerable diversity in diet, lifestyle, tooth morphology and many other ecological and evolutionary traits, the mammals included in this study all produced dental calculus deposits with a clear oral microbiome content. This suggests that the study of oral microbiome evolution and many other questions in ecology and evolution can be done using dental calculus. Second, dental calculus from hosts with herbivorous diets tends to be enriched in enzymatic inhibitors (Figure 4, Supplementary Figure 2 and 4). Therefore, special considerations must be put in place when initiating studies of herbivore dental calculus microbiome. These could include optimisations of extraction protocols that can more efficiently remove inhibitors, the use of robust DNA polymerases during library preparation, and pilot studies that allow estimating success rates for specific focal species to inform sampling strategies and sample sizes. We find that for samples with high inhibitor load using more than 10mg of dental calculus is unlikely to improve the success rate of the laboratory procedures and is unlikely to increase the proportion of unique reads in the metagenomic libraries. This result should reduce the sampling pressure on the museum specimens, preserving more of the dental calculus material for future studies. This amount of dental calculus is only a bit larger than the size of a linseed (Warinner and Fagernäs, n.d.) (Warinner and Fagernäs, n.d.). Third, although the incidence of dental calculus in animalivorous species is low (Richards et al. in review), the deposit appears to contain more DNA (Figure 3) and therefore a smaller sample amount is likely to be sufficient. These findings can guide dental calculus sampling and library preparation, facilitating the study of wild microbiomes from the past and present, while at the same time minimising the pressure of destructive sampling on museum collections.

## Acknowledgements

We would like to express our sincere gratitude to the following individuals for their invaluable contributions to this project: Frank E. Zachos, curator of the mammalian collection; Alex Bibl, collection manager; and Tim Langnitschke, scientific assistant, all from the Natural History Museum in Vienna (Austria), as well as Dominique Fonck and Mathys Rotonda from the Royal Museum for Central Africa (Belgium) for welcoming us to the collections and supporting us along the way; Gunilla Engström and E-Jean Tan from Uppsala University (Sweden) for their support during the laboratory work; Yanara Marincevic-Zunig and the rest of the team at the SNP&SEQ Technology Platform, Science for Life Laboratory, Uppsala (Sweden) for their expertise and support in sequencing (project code WI-3800); Megan Grant from the University of Edinburgh (United Kingdom) for helping gather mammal dietary data. Sequencing was performed by the SNP&SEQ Technology Platform in Uppsala. The facility is part of the National Genomics Infrastructure (NGI) Sweden and Science for Life Laboratory. The SNP&SEQ Platform is also supported by the Swedish Research Council and the Knut and Alice Wallenberg Foundation. The computations were enabled by resources provided by the National Academic Infrastructure for Supercomputing in Sweden (NAISS), partially funded by the Swedish Research Council through grant agreement no. 2022-06725. This work was supported by the Swedish Research Council (Formas) grant 2019-00275 to KG.

## Data Accessibility and Benefit Sharing

### Data Accessibility Statement

Specimen and laboratory-related metadata are available in https://github.com/markella-moraitou/mammalian_dental_calculus/tree/main/input.

Scripts used for the analysis are available in https://github.com/markella-moraitou/mammalian_dental_calculus/tree/main/scripts.

### Benefit Sharing Statement

All samples included in this study were obtained from historical specimens housed in European natural history collections. Sample collection and processing methodologies were approved by the respective curators prior to the commencement of the study.

### Author Contributions

KG, MM and JR conceived the study. MM and JR collected samples and performed laboratory work. MM analysed the data and produced figures, with contribution from CB. FEZ, KS, EG, ZT, ACK, OP, RS, PK, and RPM provided logistical support and access to samples and curated sample metadata. MM wrote the manuscript with input from KG and all co-authors. KG administered the project, acquired funding and resources.

