## Supplementary Figures for "Host traits impact the outcome of metagenomic library preparation from dental calculus samples across diverse mammals"

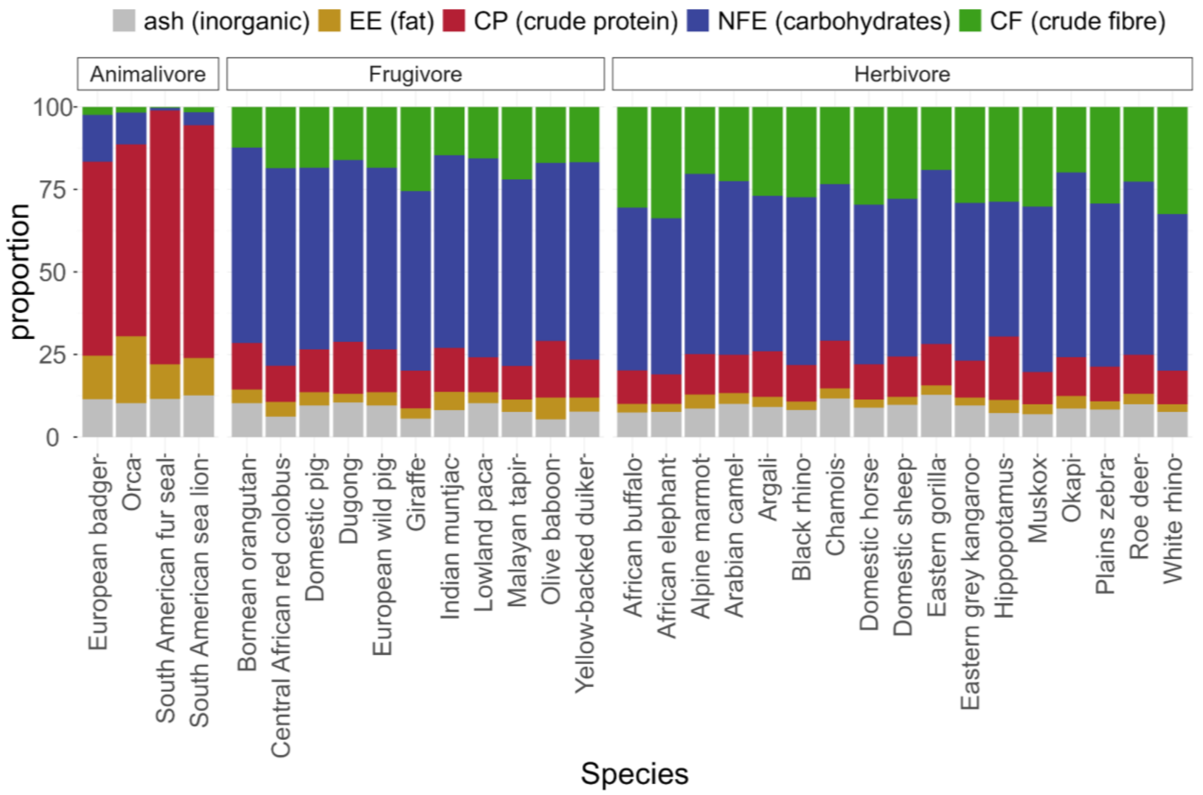


**Supplementary Figure 1.** Barplot showing chemical composition of the diet of study species according to Lintulaakso et al. (2023) (data in Supplementary Table 2).

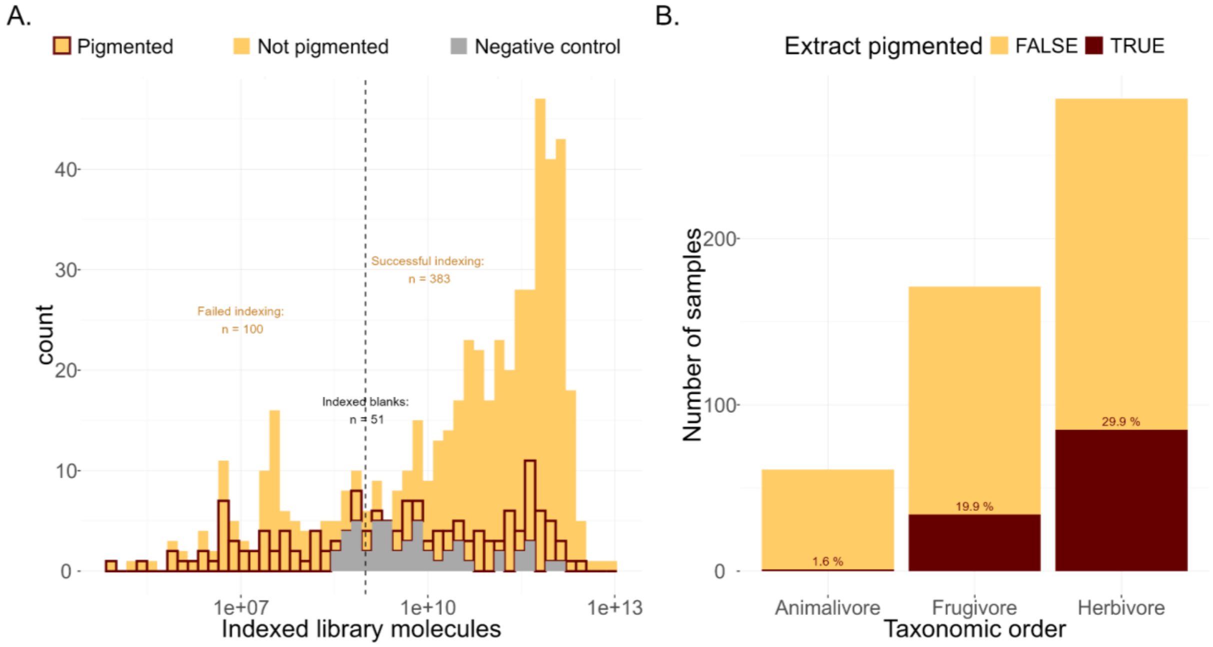


**Supplementary Figure 2.** A) Distribution of indexed library molecules for samples (yellow) and blanks (grey) (note the log10 scale on the x-axis). Brown outlines show libraries from pigmented extracts. The vertical dashed line at $10^9$ indexing copies was used as a cut-off for a successful indexed library and corresponds to ca. the first quartile of indexed library molecules for the negative controls ($1.2x10^9$). B) Proportion of samples in our dataset that generated a pigmented DNA extract, per dietary category, indicating possible co-elution of inhibitors.

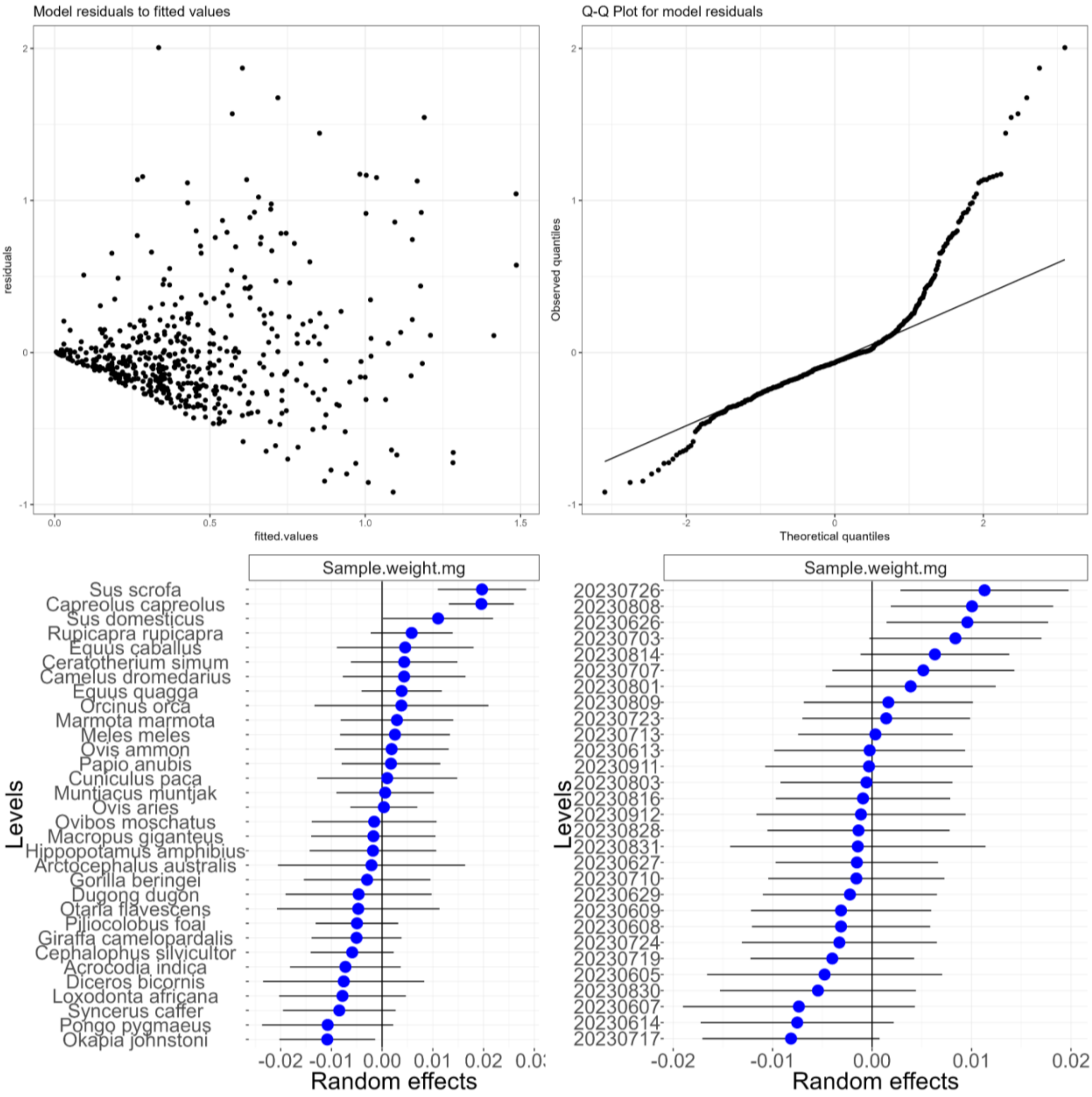


**Supplementary Figure 3.** Diagnostic plots and random effects for Model 1. Top left: Plot of residuals to fitted values. Top right: Quantile-Quantile (QQ) plot for model residuals. Bottom: left: Random effects for mammalian host species. Bottom right: Random effects for DNA extraction batch.

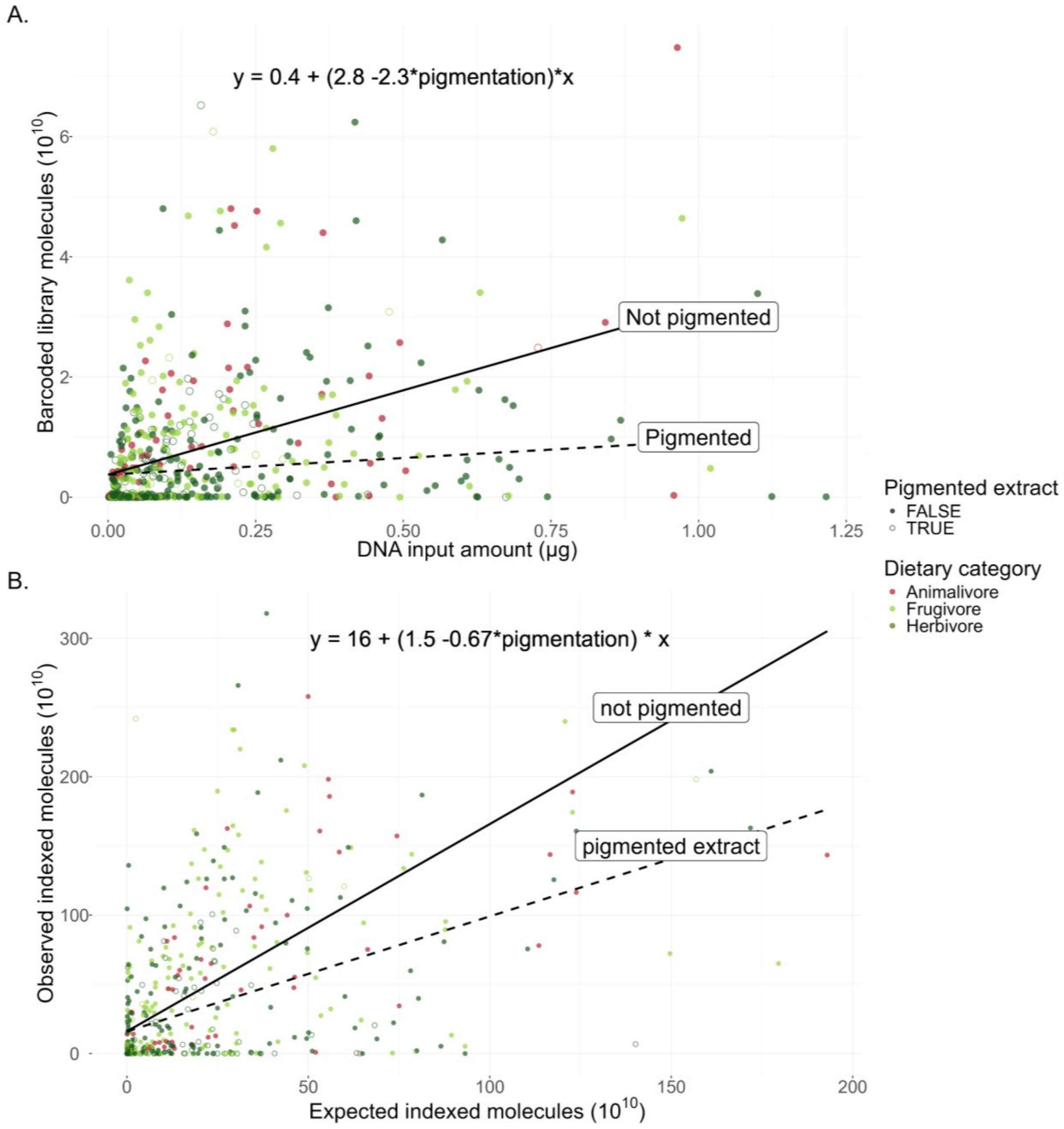


**Supplementary Figure 4.** A) The relationship between DNA input in µg and barcoded library output for 497 samples that underwent barcoding, and B) the relationship between expected and observed indexed library output for 472 samples that underwent indexing. The formulas on the top are based on all significant model estimators (in both cases, those are the intercept, the input and extract pigmentation) and the two lines show the fitted values for a samples with pigmented (dashed) or nonpigmented (solid) DNA extracts according to a linear mixed model.

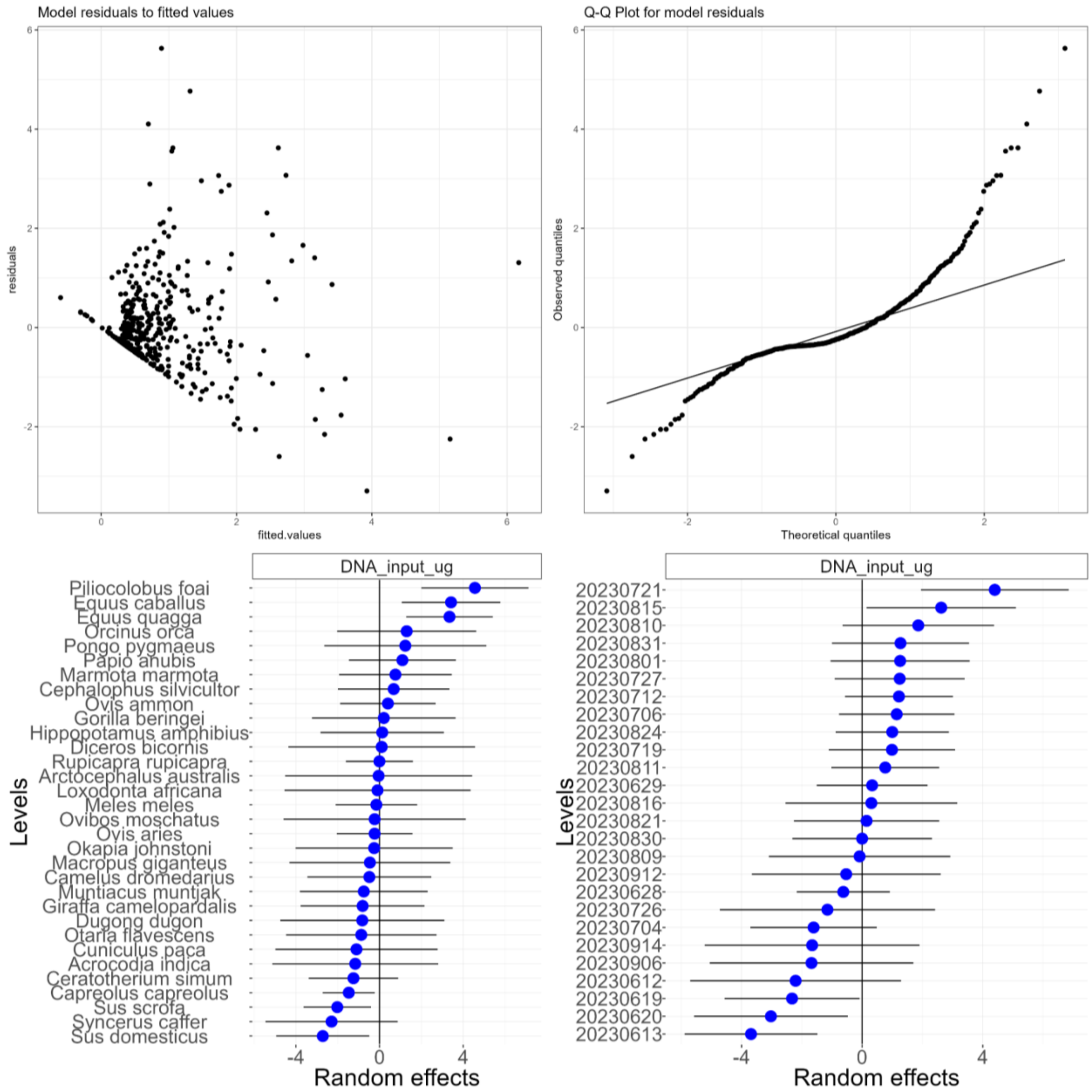


**Supplementary Figure 5.** Diagnostic plots and random effects for Model 2. Top left: Plot of residuals to fitted values. Top right: Quantile-Quantile (QQ) plot for model residuals. Bottom: left: Random effects for mammalian host species. Bottom right: Random effects for adapter ligation batch.

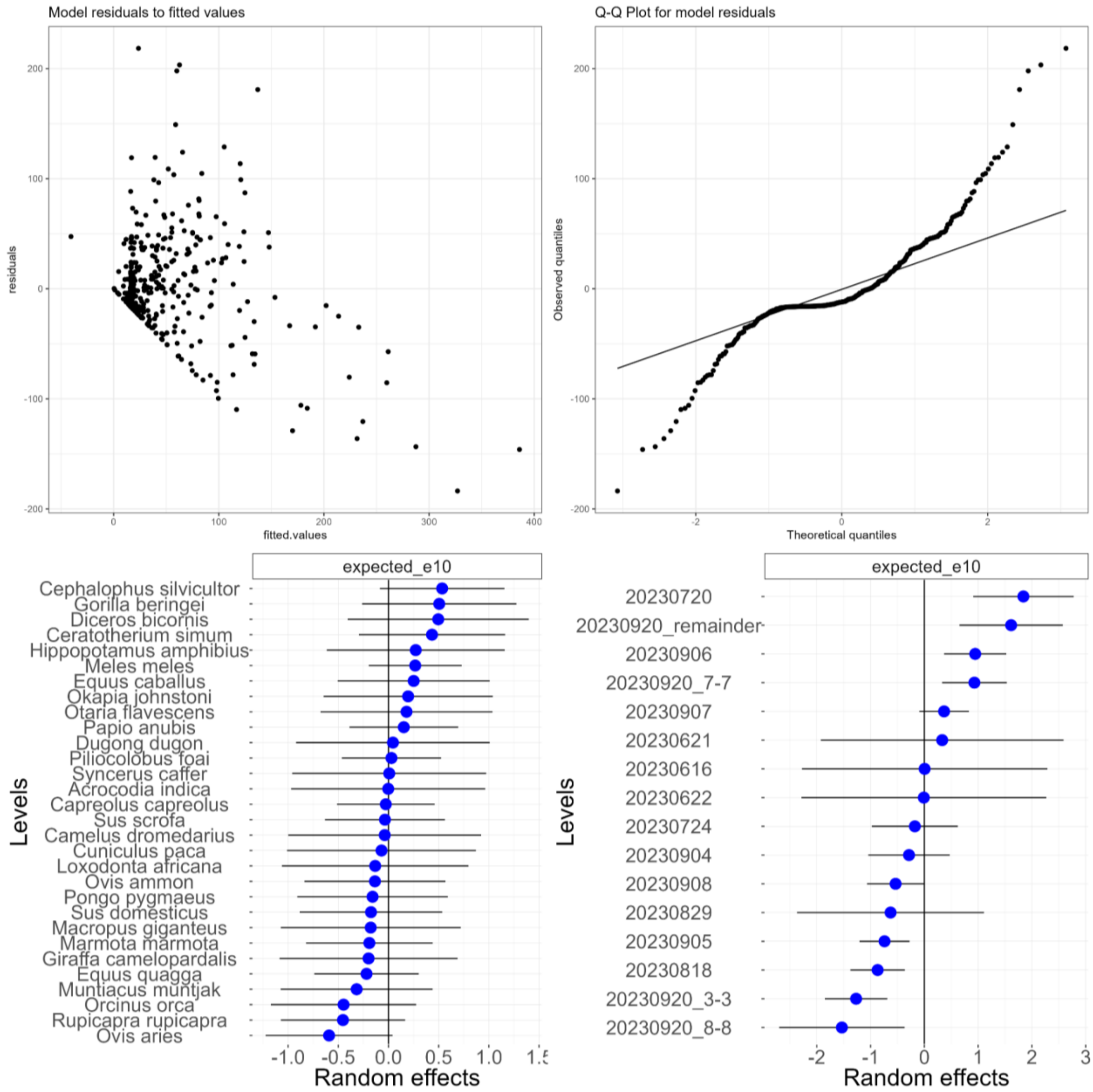


**Supplementary Figure 6.** Diagnostic plots and random effects for Model 3. Top left: Plot of residuals to fitted values. Top right: Quantile-Quantile (QQ) plot for model residuals. Bottom: left: Random effects for mammalian host species. Bottom right: Random effects for indexing batch.

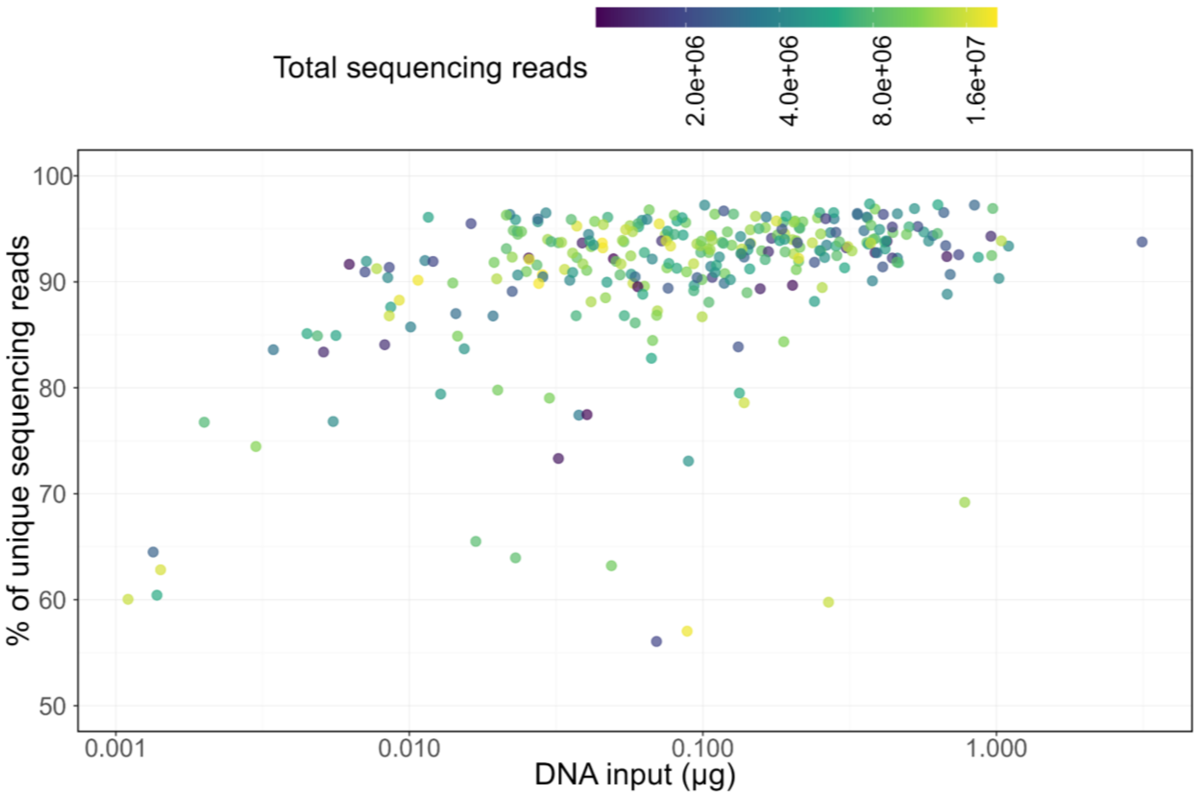


**Supplementary Figure 7.** Percentage of unique sequencing reads plotted against the initial DNA input (in µg) used for library preparation. The colour indicates the total sequencing depth for each sample. For this visualisation, we excluded samples with fewer than $10^6$ or more than $2×10^7$ reads; The remaining 304 samples are presented here. Note the log10-transformed x-axis and colour scale.

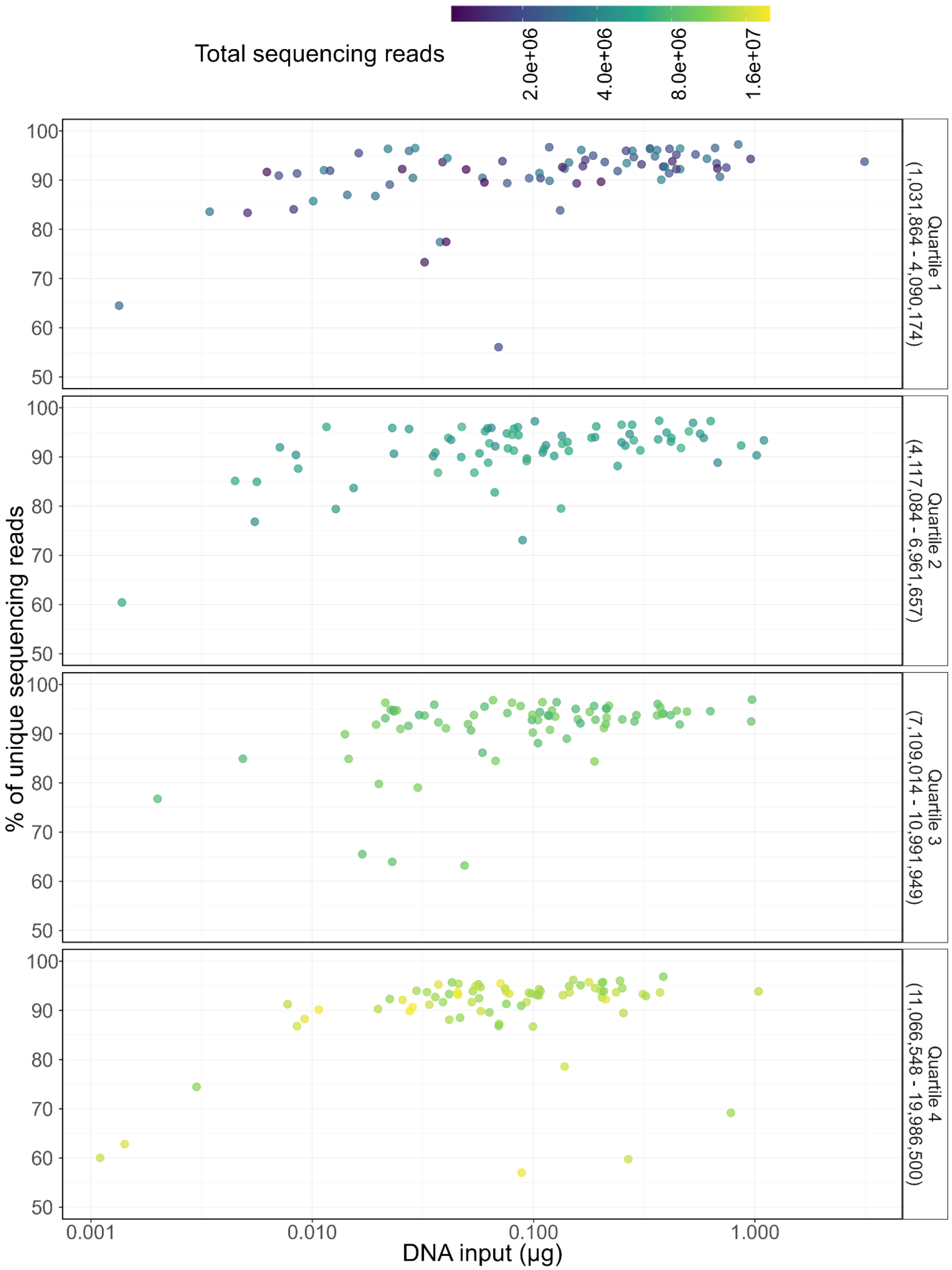


**Supplementary Figure 8.** Percentage of unique sequencing reads plotted against the initial DNA input (in μg) used for library preparation presented separately for each quartile of sequencing depth. Like in Supplementary Figure 10, this visualisation includes 304 samples with sequencing depth between $10^6$ and $2{\times}10^7$ reads. Note the log10-transformed x-axis and colour scale.

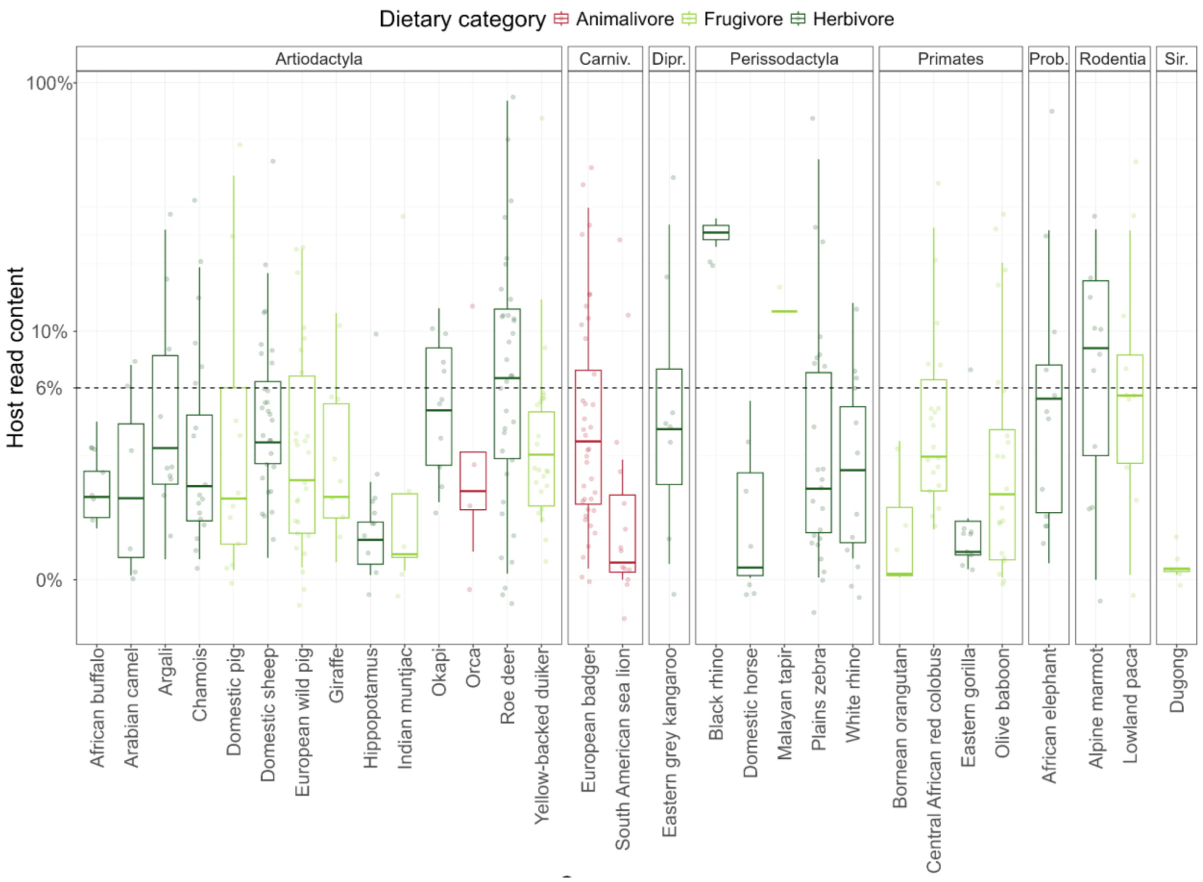


**Supplementary Figure 9.** Percentage of host reads in dental calculus for different species. Colour indicates dietary category based on Lintulaakso et al. (2023). Facets reflect different taxonomic orders. Note the log10-transformed y axis to increase the visibility of low values. Abbreviations: Carniv. = Carnivora, Dipr. = Diprotodontia, Prob. = Proboscidea, Sir. = Sirenia. The horizontal dashed line indicates the average host read percentage (5.9%).

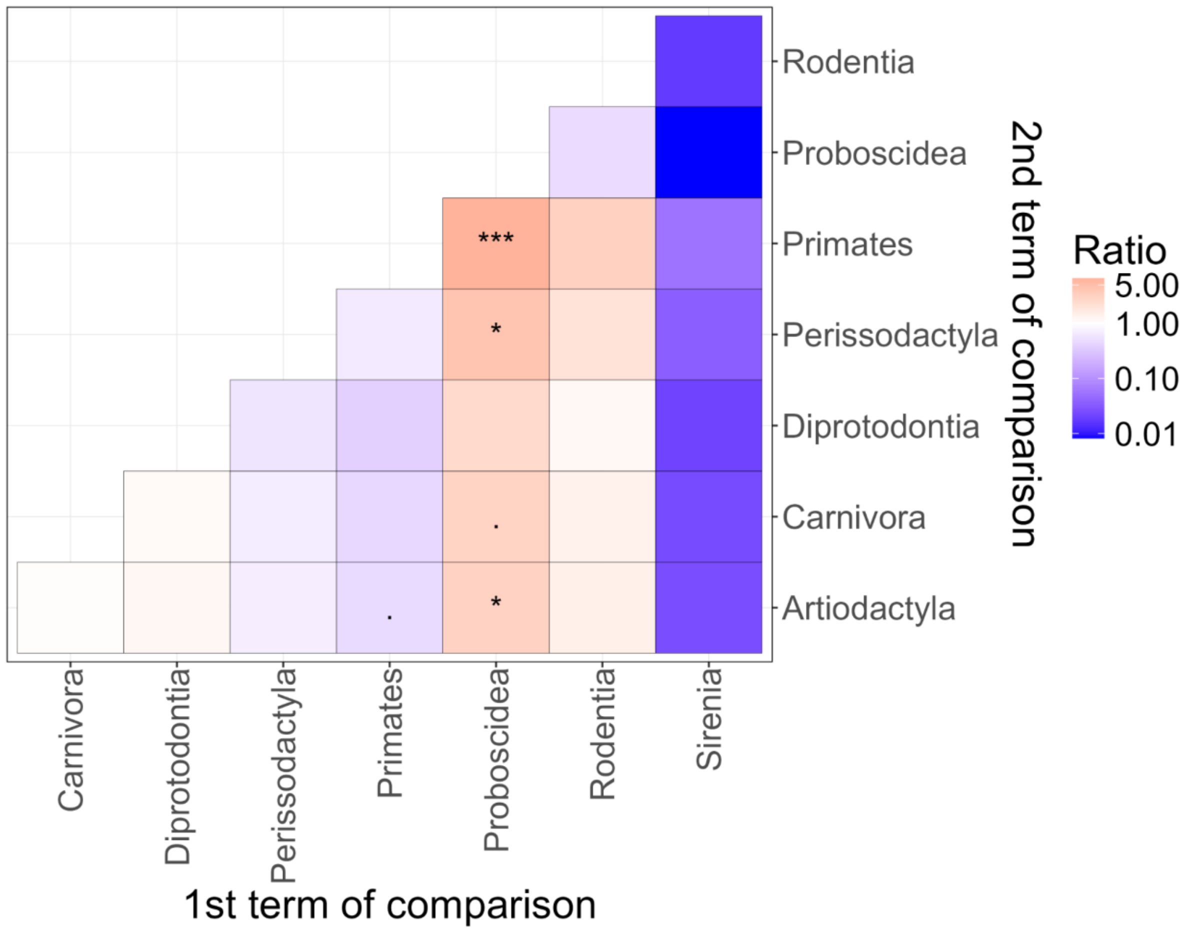


**Supplementary Figure 10.** Ratio of host DNA content (%) between orders. Red indicates that the host taxonomic order on the x-axis has a larger proportion of host reads than the order on the y-axis, while blue indicates the reverse. Significance levels: (***) $p < 0.001$, (**) $p < 0.01$, (*) $p < 0.05$, (.) $p < 0.1$.

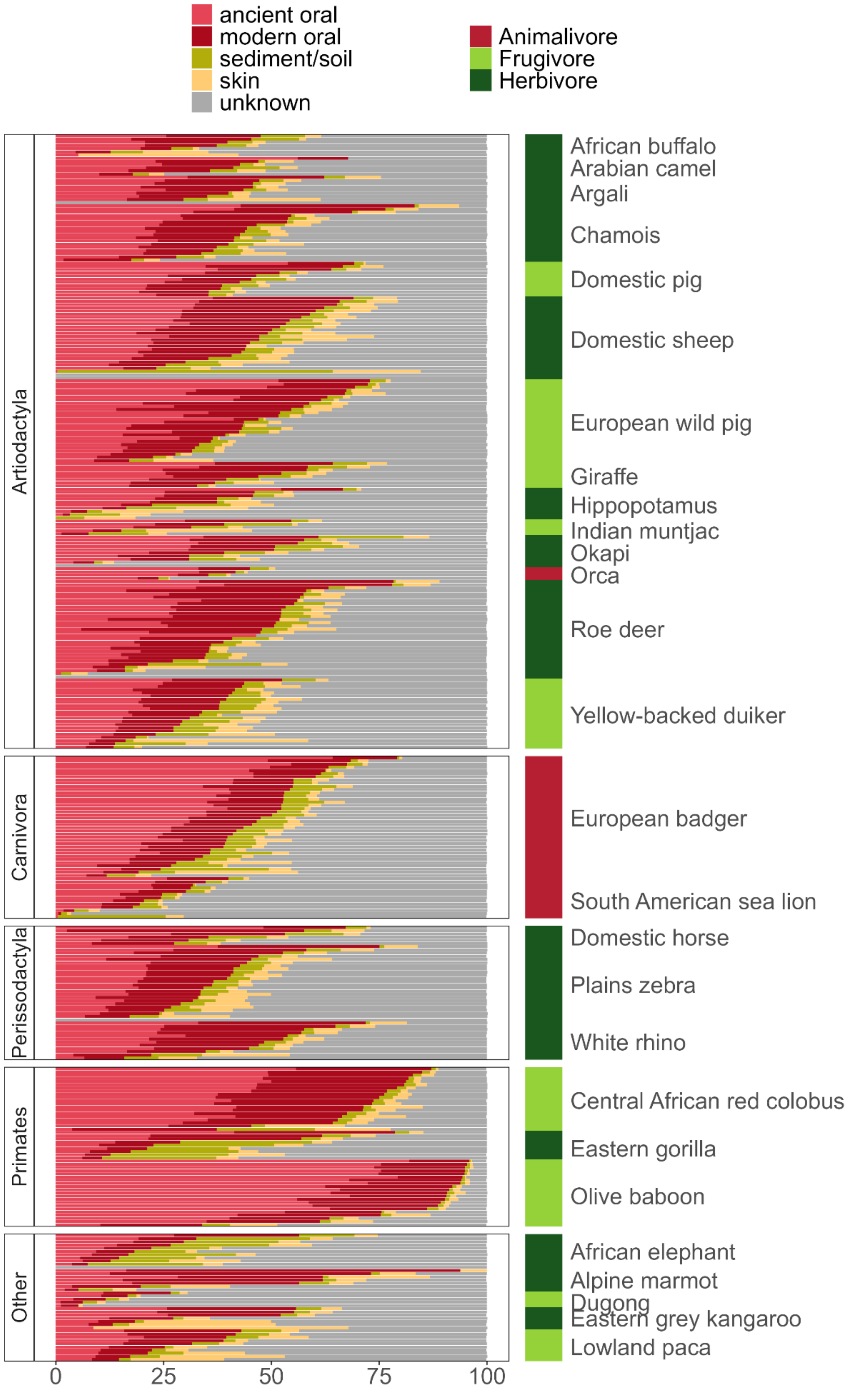
ancient oral
modern oral
sediment/soil
skin
unknown
Animalivore
Frugivore
Herbivore
Artiodactyla
Carnivora
Perissodactyla
Primates
Other
African buffalo
Arabian camel
Argali
Chamois
Domestic pig
Domestic sheep
European wild pig
Giraffe
Hippopotamus
Indian muntjac
Okapi
Orca
Roe deer
Yellow-backed duiker
European badger
South American sea lion
Domestic horse
Plains zebra
White rhino
Central African red colobus
Eastern gorilla
Olive baboon
African elephant
Alpine marmot
Dugong
Eastern grey kangaroo
Lowland paca
0
25
50
75
100

**Supplementary Figure 11.** Estimated composition of dental calculus metagenomes (after removing reads mapping to the host and human genome) with regard to four types of microbiome sources: human ancient oral, human modern oral, human skin and sediment/soil. The proportions were estimated using a kmer-based approach implemented in decOM (Duitama González et al. 2023). The “Unknown” partition refers to sequences that were found in the samples but none of the sources, however it's a feature under development and may not be entirely accurate. The bar on the right side indicates the dietary category of the mammalian host. Plot data can be found in Supplementary Table 1.
